# Dub-seq: dual-barcoded shotgun expression library sequencing for high-throughput characterization of functional traits

**DOI:** 10.1101/387399

**Authors:** Vivek K. Mutalik, Pavel S. Novichkov, Morgan N. Price, Trenton K. Owens, Mark Callaghan, Sean Carim, Adam M. Deutschbauer, Adam P. Arkin

## Abstract

A major challenge in genomics is the knowledge gap between sequence and its encoded function. Gain-of-function methods based on gene overexpression are attractive avenues for phenotype-based functional screens, but are not easily applied in high-throughput across many experimental conditions. Here, we present <u>**Du**</u>al <u>**B**</u>arcoded <u>**S**</u>hotgun <u>**E**</u>xpression Library Se<u>**q**</u>uencing (Dub-seq), a method that greatly increases the throughput of genome-wide overexpression assays. In Dub-seq, a shotgun expression library is cloned between dual random DNA barcodes and the precise breakpoints of DNA fragments are associated to the barcode sequences prior to performing assays. To assess the fitness of individual strains carrying these plasmids, we use DNA barcode sequencing (BarSeq), which is amenable to large-scale sample multiplexing. As a demonstration of this approach, we constructed a Dub-seq library with total *Escherichia coli* genomic DNA, performed 155 genome-wide fitness assays in 52 experimental conditions, and identified 813 genes with high-confidence overexpression phenotypes across 4,151 genes assayed. We show that Dub-seq data is reproducible, accurately recapitulates known biology, and identifies hundreds of novel gain-of-function phenotypes for *E. coli* genes, a subset of which we verified with assays of individual strains. Dub-seq provides complementary information to loss-of-function approaches such as transposon site sequencing or CRISPRi and will facilitate rapid and systematic functional characterization of microbial genomes.

**Importance:** Measuring the phenotypic consequences of overexpressing genes is a classic genetic approach for understanding protein function; for identifying drug targets, antibiotic and metal resistance mechanisms; and for optimizing strains for metabolic engineering. In microorganisms, these gain-of-function assays are typically done using laborious protocols with individually archived strains or in low-throughput following qualitative selection for a phenotype of interest, such as antibiotic resistance. However, many microbial genes are poorly characterized and the importance of a given gene may only be apparent under certain conditions. Therefore, more scalable approaches for gain-of-function assays are needed. Here, we present **Du**al **B**arcoded **S**hotgun **E**xpression Library Se**q**uencing (Dub-seq), a strategy that couples systematic gene overexpression with DNA barcode sequencing for large-scale interrogation of gene fitness under many experimental conditions at low cost. Dub-seq can be applied to many microorganisms and is a valuable new tool for large-scale gene function characterization.

## INTRODUCTION

Advances in DNA sequencing have had a tremendous impact on microbial genomics, as thousands of genomes have now been sequenced^1^. However, only a small fraction of these microorganisms have been experimentally studied and as such, our predictions of gene function, metabolic capability, and community function for these microorganisms are based largely on automated computational approaches^2^. Unfortunately, many of these computational predictions are incomplete or erroneous, especially in instances where the homology of a sequenced gene is too distant from any experimentally characterized relative^3^. To bridge this gap between sequencing and functional characterization, it is imperative that large-scale, inexpensive, and organism-agnostic tools are developed and applied^4^.

A number of large-scale approaches based on loss-of-function genetics have been developed for microorganisms including gene-knockout libraries^5-9^, recombineering based methods^10,11^, transposon mutagenesis coupled to next-generation sequencing (TnSeq)^12,13^, and CRISPR interference (CRISPRi)^14^. Collectively, these strategies all rely on measuring the phenotypic consequences of removing a gene from a microorganism and inferring protein function based on these phenotypes. An adaptation of TnSeq that incorporates and uses random DNA barcodes (RB-TnSeq) to measure strain abundance in a competitive growth assay^13^ has recently been applied on a larger scale to identify mutant phenotypes for thousands of genes across 32 bacteria^15^. Despite their utility, these loss-of-function approaches suffer some limitations: only CRISPRi is effective for interrogating essential genes under multiple conditions, it is challenging to identify phenotypes for genes with redundant functions using single mutants, and these approaches require some degree of genetic tractability in the target microorganism.

A complimentary approach for studying gene and organism function is to generate gain-of-function overexpression libraries and analyze the phenotypic consequences of increased gene dosage. Indeed, the impact of enhanced gene dosage on adaptation and evolution are well documented across all three kingdoms of life and have been shown to be an important contributor to numerous diseases and drug-resistance phenotypes^16-18^. Overexpression as a genetic tool has a rich history of connecting genes to cellular functions and has been exploited as a versatile screening technique to identify drug targets^16,19,20^, antibiotic and metal resistance genes^17,21,22^, virus-resistance genes^23^, genetic suppressors^24,25^, as well as for a number of chemical genomics^8,9^ and biotechnology applications^26-28^. While a number of technologies have been developed for overexpression screens including defined open reading frame (ORF) libraries^6,20,29^ and activation modes of recombineering^30,31^, transposon insertions^32^ or CRISPR systems^33^, these strategies are limited, either due to the need for expensive and laborious generation of archived strains or the need for organism-specific genetic tools.

A simpler alternative for overexpression screens is a shotgun library-based approach in which random DNA is introduced into a host organism for phenotyping and functional assessment. This approach has been widely used for studying increased-copy number effects on a desired phenotype^26,27^ and for activity-based screening of metagenomic samples^34,35^. Nevertheless, most shotgun expression libraries have only been assayed in a small number of conditions looking for a specific gene-function, and are often performed as qualitative selections on a plate^34-36^. Furthermore, current shotgun-based approaches typically require tedious and expensive sequencing and sample preparation protocols for identifying the selected gene(s)^26,27,37,38^. With arrival of next-generation sequencing technologies, all positive candidates can be pooled, and cloned regions can be amplified and sequenced in parallel^39,40^. Unfortunately, sequencing the cloned regions (to identify the genes conferring the phenotype) is labor intensive and may become cost-prohibitive if the overexpression library is being assayed in many conditions. As such, there is a need for high-throughput gain-of-function technology that is simple, quantitative, agnostic to source DNA, and which facilitates multiplexed quantification of fitness under hundreds of experimental conditions.

Here we present a new method termed Dub-seq, or dual barcoded shotgun expression library sequencing, for performing high-throughput and quantitative gain-of-function screens. Dub-seq requires an initial characterization of the overexpression library by linking the genomic breakpoints of each clone to a pair of random DNA barcodes. Subsequent screens are performed using a competitive fitness assay with a simple DNA barcode sequencing and quantification assay (BarSeq^41^). As a demonstration of this approach, we generated an *E. coli* Dub-seq library and assayed the phenotypic consequences of overexpressing nearly all genes on *E. coli* fitness under dozens of experimental conditions. We show that Dub-seq yields gene fitness data that is consistent with known biology and also provides novel gene-function insights. We validate some of these new findings by overexpressing individual genes and quantifying these strains’ fitness. Given that only DNA and a suitable host organism for assaying fitness are necessary, Dub-seq can be readily extended to diverse functional genomics and biotechnology applications.

## RESULTS

### Overview of Dub-seq

The Dub-seq approach is summarized in **Figure 1** and can separated into four different steps. First, a plasmid library is generated with pairs of random 20 nucleotide DNA sequences, termed the UP and DOWN barcodes. To link the identities of the two-barcode sequences on each plasmid, <u>B</u>arcode-<u>P</u>air <u>seq</u>uencing (BPseq) is performed (**Fig. 1a**, Methods). Second, sheared genomic DNA from an organism under investigation is cloned between the previously associated UP and DOWN barcodes (**Fig. 1b**). Third, the genomic fragment endpoints are mapped and associated with the two-barcode sequences using a TnSeq-like protocol^13^. We term this step <u>B</u>arcode-<u>A</u>ssociation-with <u>G</u>enome fragment by <u>seq</u>uencing or BAGseq and the resulting plasmid library as the “Dub-seq” library (**Fig. 1c**). The BAGseq step requires two sample preparations to separately map genomic fragment junctions to the UP and DOWN barcodes. The BAGseq characterization generates a table of barcode sequences and the cloned chromosomal breakpoints at single-nucleotide resolution. Because the two random DNA barcodes have been previously associated, we can infer the exact sequence of each plasmid in the Dub-seq library if the sequence of the source DNA is known. Lastly, we introduce the Dub-seq plasmid library into a host bacterium and monitor the fitness of strains carrying these plasmids in a competitive fitness assay under a particular condition by PCR amplifying and quantifying the abundance of the DNA barcode sequences (BarSeq^41^, **Fig. 1d**). In these pooled fitness experiments, the barcode abundance changes depending upon the fitness phenotype imparted by the barcode-associated-genome fragments. A data analysis pipeline yields fitness scores for individual strains (or “fragments”) and for each gene. These gene scores provide an assessment of the phenotypic consequence of overexpressing nearly all of the genes represented in the cloned DNA fragments. The advantage of Dub-seq is that it decouples the characterization of a shotgun overexpression library (which is more laborious) from the cheaper and simpler fitness determination step using BarSeq. As such, a Dub-seq library can be readily assayed in hundreds of different experimental conditions. Dub-seq can be viewed as an overexpression-based, gain-of-function version of our previously described method for random barcode transposon-site sequencing (RB-TnSeq)^13^.

**Figure 1.**
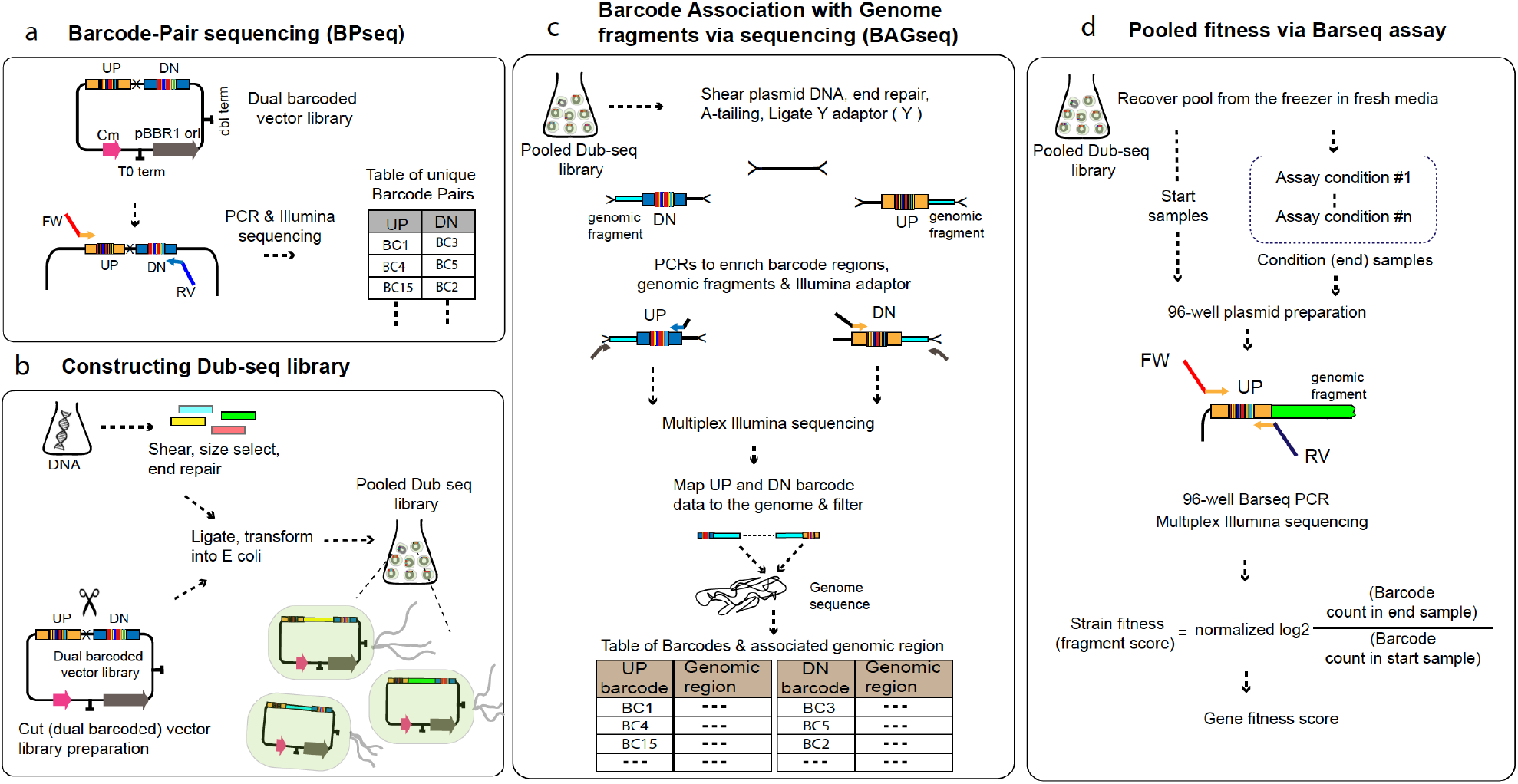
Schematic overview of the Dub-seq approach. (a) A pair of random 20 nucleotide DNA sequences, the UP and DOWN (DN) barcodes are cloned into an expression vector. Deep sequencing of the dual barcoded vector (BPseq) associates UP and DOWN barcode sequences. (b) Target genomic DNA is randomly sheared and cloned between the UP and DOWN barcodes to create the Dub-seq plasmid library. (c) To characterize the Dub-seq library, a “Tn-seq” like protocol is performed to precisely map the two genomic breakpoints of each insert and to associate each breakpoint with its random DNA barcode sequence. If the source genome(s) has been sequenced, then BAGseq can be used to define the exact sequence of each plasmid in the library. (d) The fitness of bacteria carrying different plasmids can be measured with pooled growth assays and deep sequencing of the DNA barcodes (BarSeq). Strain (or fragment) fitness is defined as the log2 ratio of barcode abundance after selection (end) versus before (start). Gene fitness is estimated from the fragments’ fitness by a constrained regression.

### Generation of *E. coli* Dub-seq library

To generate a Dub-seq library, we used a broad host range vector with a pBBR1 replication origin. We used standard molecular biology techniques to insert two random 20 nucleotide barcode sequences on the plasmid, the UP and DOWN barcodes, that juxtapose a unique Pmil restriction enzyme site on the plasmid. Both the UP barcodes and DOWN barcodes contain common PCR priming sites for rapid amplification of all barcodes from a pooled sample. We generated a dual barcoded vector library with ~250,000 clones in *E. coli* and characterized this library by associating the barcode pairs using BPseq. The vector library of ~250,000 clones was sufficient to map unique barcode-pairs with confidence and also to yield a Dub-seq library in which each fragment will have a unique barcode (see below).

To generate the *E. coli* Dub-seq library, we extracted *E. coli* (BW25113) genomic DNA, sheared to 3 kb fragment size, and cloned the fragments into the dual barcoded backbone vector digested with PmiI. The *E. coli* Dub-seq library encompasses ~40,000 vectors, corresponding to about 8X coverage of the *E. coli* genome. In this study, we used the endogenous *E. coli* transcription and translation apparatus to drive the expression of the encoded gene(s) within each genomic fragment, although future studies could use inducible systems (for example, when the source of the cloned Dub-seq DNA differs from the host bacterium for assaying fitness^42^).

We next characterized the *E. coli* Dub-seq library using BAGseq, which identifies the cloned genome fragment and its pairings with the neighboring dual barcodes. As there are two barcodes for each Dub-seq library, we performed two separate BAGseq sample preparation steps, one for the UP barcodes and one for the DOWN barcodes. Briefly, BAGseq involves shearing of the Dub-seq plasmid library, end repair, Illumina adaptor ligation, PCR amplification of the junction between the barcode and genomic insert using primers that are complementary to one of the barcode-specific primer binding sites, and deep sequencing of these samples (modified from reference 11). After filtering out barcodes that mapped to more than one genomic fragment, we identified 30,558 unique barcode pairs that we could confidently associate with a genomic fragment.

In the *E. coli* Dub-seq library, the fragments are evenly distributed across the chromosome (**Fig. 2a**), the average fragment size is 2.6 kB (**Fig. 2b**), and the majority of fragments covered 2-3 genes in their entirety (**Fig. 2c**). 80% of genes in the *E. coli* genome are covered (from start to stop codon) by at least 5 independent genomic fragments in the Dub-seq library (**Fig. 2d**) and 97% of all genes are covered by at least one fragment. Just 135 genes are not covered in their entirety by any Dub-seq fragment (**Supplementary Table 1**). Many of these unmapped or uncovered genes encode membrane and ribosomal proteins and probably reflect the lethality of overexpressing these genes^43^. Other genes could not be confidently mapped because they are associated with repetitive regions. For example, we could not confidently map fragments covering ETT2 type III secretion system pathogenicity island and its regulator gene *ygeH* which has tetratricopeptide repeat motifs, while the neighboring protein-coding genes are well mapped (**Fig. 2a**). Similarly, we could not map genes within ribosomal RNA operons (example, *rrlD*, **Fig. 2a**), as *E. coli* encodes multiple nearly-identical copies of these loci. Some large genes with length more than 3.5 Kb, such as *rpoB*, are not entirely covered by any fragments in our library, while other large genes such as *acrB* are covered by only one fragment (**Fig. 2a**).

**Figure 2.**
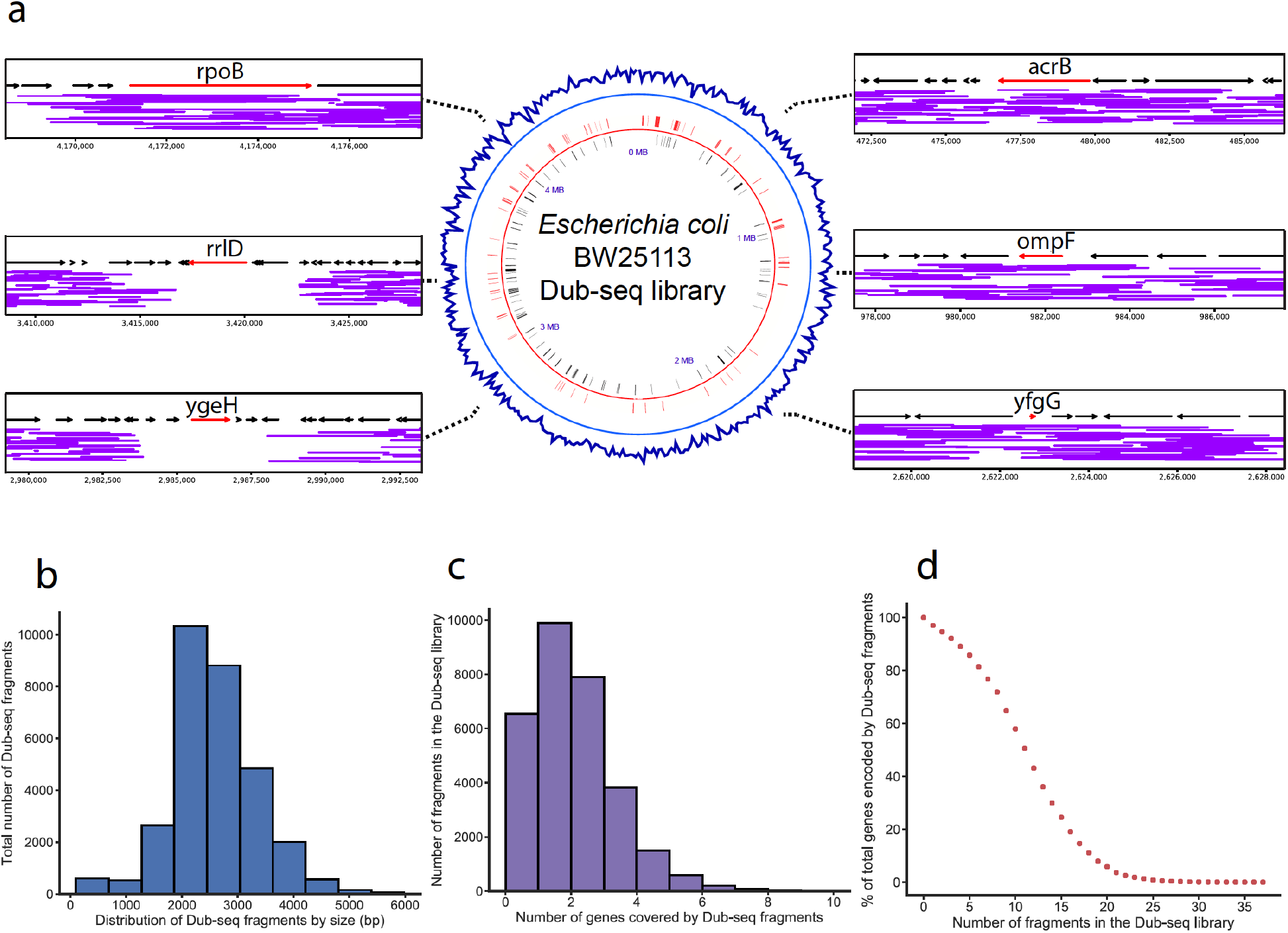
*E. coli* Dub-seq library characterization. (a) Center: genomic coverage of the *E. coli* BW25113 Dub-seq library in 10 kB windows (blue track). Black and red line-tracks represent genes essential for viability when deleted^5^ that are encoded on the negative and positive strands, respectively and are covered in the Dub-seq library. Left and right: regions of the *E. coli* chromosome covering *acrB, ompF, yfgG, ygeH, rrlD* and *rpoB.* Each purple line represents a Dub-seq genomic fragment(the y-axis is random). (b) The fragment insert size distribution in the *E. coli* Dub-seq library. (c) The distribution of number of genes that are completely covered (start to stop codon) per genomic fragment in the *E. coli* Dub-seq library. (d) Cumulative distribution plot showing the percentage of genes in the *E. coli* genome (y-axis) covered by a number of independent genomic fragments (x-axis).

Of the *E. coli* protein-coding genes that are essential for viability when deleted^5^, 95% are completely covered by at least one fragment in the Dub-seq library (**Supplementary Table 2**). This demonstrates that the Dub-seq approach can interrogate genes that are not typically assayed for conditional phenotypes in loss-of-function approaches. There are only 17 protein-coding genes that are both essential for viability when deleted and absent from our Dub-seq library (**Supplementary Table 2**).

### Strain and gene fitness profiling using BarSeq

The key advantage of Dub-seq is the ease of assessing the relative fitness contributions of all genes contained in the cloned genomic fragments using pooled, competitive growth assays. Depending on the assay condition and the gene(s) encoded by a genomic fragment, the relative abundance of a strain carrying that fragment can change due to its fitness advantage or disadvantage relative to strains carrying other fragments. Because the DNA barcodes have been previously associated to each genomic fragment, we can simply compare the relative abundance of each barcode before and after selective growth using DNA barcode sequencing or BarSeq^41^.

As a demonstration of Dub-seq fitness assays and to illustrate our approach for calculating strain (fragment) and gene fitness scores, we recovered an aliquot of the *E. coli* Dub-seq library in LB to mid-log phase, collected a cell pellet for the “start” (or time-zero sample), and used the remaining cells to inoculate an LB culture supplemented with 1.2 mM nickel. After growth in the presence of nickel, we collected a second cell pellet for the “condition” sample. We extracted plasmid DNA from the start and condition samples, PCR amplified the UP and DOWN DNA barcodes from each, and sequenced the DNA barcodes with Illumina. We calculate the fragment fitness score for each strain by taking the normalized log2 ratio of the number of reads for each barcode in condition sample versus the start sample (**Fig. 1**). Positive scores indicate that the gene(s) contained on that fragment lead to an increase in relative fitness, while negative values mean the gene(s) on the fragment reduced relative fitness. Scores near zero indicate no fitness reduction or benefit for the gene(s) under the assayed condition. As in previous work^44^, we find that fitness scores calculated with either UP barcodes or DOWN barcodes yield very similar results (*r* = 0.94, **Supplementary Fig. 1ab**). Therefore, we only sequenced the UP barcodes for all additional experiments in this study.

Given that multiple, causative and non-causative genes can be contained on a single fragment, to assign a fitness score to a particular gene it is necessary to examine the score of all fragments containing the gene. Here, we considered two different ways to estimate fitness score of a gene. The first approach was to simply take the average of all fitness scores for fragments that contained the gene in its entirety (the “mean” score). The second approach was to use a regression method for estimating gene fitness score so as to prevent genes from having artifactually high fitness scores if they were located near other causative genes. Specifically, we adopted non-negative least squares regression (the “regression” score) (see Methods). To illustrate how the mean and regression scores differ in practice, consider the gene fitness scores for two adjacent genes under elevated nickel stress, *rcnA* and *rcnR* (**Fig. 3a and 3b**). RcnA is a nickel efflux protein whose overexpression is known to lead to increased nickel tolerance^45^. Conversely, *rcnR* encodes a transcriptional repressor that weakly represses its own expression and that of *rcnA*, and the overexpression of *rcnR* alone is not expected to increase nickel tolerance^45^. While the mean and regression approaches both result in similar (and correct) high Dub-seq scores for *rcnA* (**Fig. 3a**), only the regression approach results in the correct, neutral fitness score for the *rcnR* (**Fig. 3b**). The mean score calculation approach leads to an artifactually high fitness score for *rcnR* because many of the fragments that contain this gene also contain the neighboring *rcnA* (**Fig. 3b, Supplementary Figs. 2ab and 3ab**). Based on these results and other examples (**Supplementary Fig. 4**) that we examined, we concluded that the optimal strategy was to use the regression method for calculating Dub-seq gene fitness scores (Methods).

**Figure 3.**
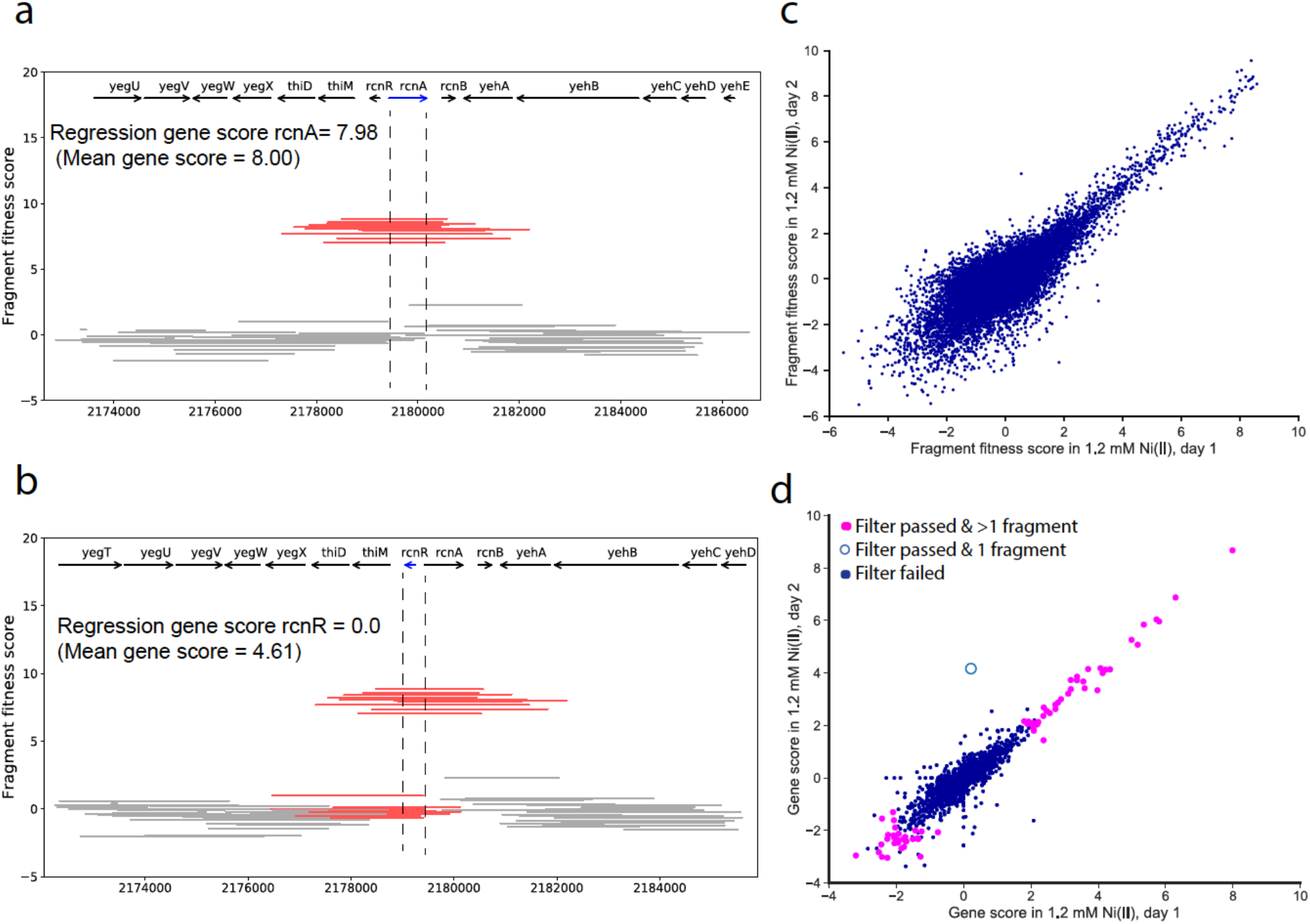
Fragment and gene fitness Dub-seq scores. (a) Dub-seq fragment (strain) data for region surrounding *rcnA* under elevated nickel stress (y-axis). Each line shows a Dub-seq fragment. Those that completely cover *rcnA* are in red. Both the mean and regression scores reflect the known biology of *rcnA* as a nickel resistance determinant^45^. (b) Same as (a) for the neighboring *rcnR*, which encodes a transcriptional repressor of *rcnA.* Fragments that cover *rcnR* are in red. (c) Comparison of fragment fitness scores for two biological replicates of 1.2 mM nickel stress. (d) Same as (c) for gene fitness scores calculated using the regression approach. Genes are highlighted if their data passed our statistical filters for reliable effects (see Methods); we also show whether the gene score is based on just one fragment.

To assess the reproducibility of Dub-seq fitness assays, we compared the results obtained from independent samples. First, the number of sequencing read counts for each UP barcodes from the Dub-seq library from different start samples were highly correlated (**Supplementary Fig. 1c**). Likewise, between two biological replicates of the nickel stress experiment, we found a strong correlation for fragment fitness (*r* = 0.80; **Fig. 3c**) and for regression-based gene fitness (*r* =0.89; **Fig. 3d**).

### Fitness profiling across dozens of experimental conditions

To demonstrate the scalability of Dub-seq, we performed 155 genome-wide pooled fitness experiments representing 52 different chemicals: 23 compounds as the sole source of carbon in a defined growth media and varying concentrations of 29 inhibitory compounds in rich media (**Fig. 4**). The inhibitory compounds included metals, salts, and antibiotics. For each of these assays, we compared the abundance of the UP barcodes before and after growth selection. We multiplexed 48 or 96 BarSeq PCR samples per lane of Illumina sequencing, at a sequencing cost of about $20 per genome-wide assay. In the typical condition sample, we obtained ~4.2 million BarSeq reads, representing ~100 reads on an average for each clone in the Dub-seq plasmid library. We computed gene fitness scores (using the regression approach) for 4,027 protein-coding genes and for 124 RNA genes. The gene fitness scores were reproducible, with a median pairwise correlation of 0.80 across 64 biological replicates.

**Figure 4.**
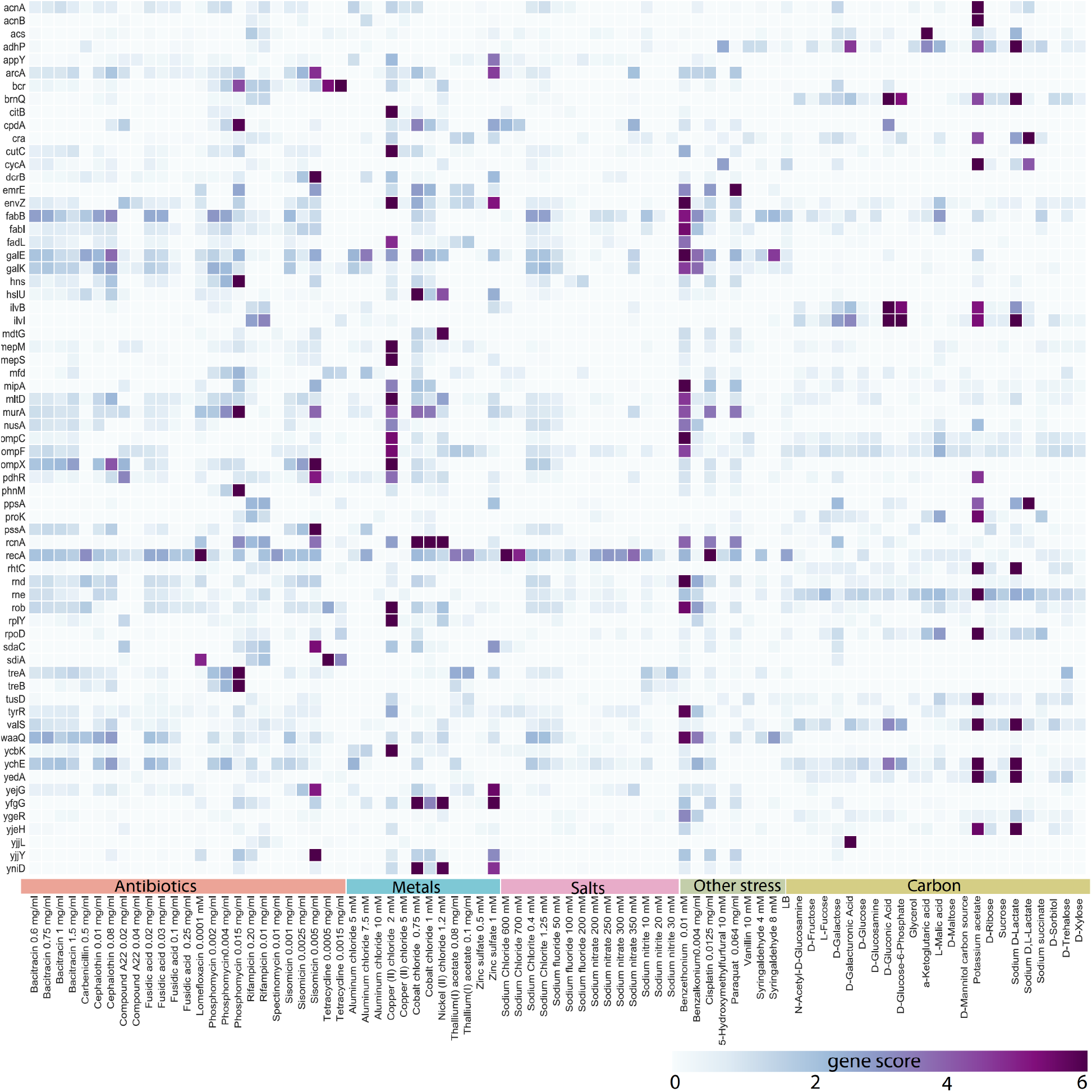
Heatmap of Dub-seq fitness data for 53 conditions and for 67 genes with large benefits. Only genes with a high-confidence effect and gene fitness score >= 6 in at least one condition are shown. Gene scores from replicate experiments were averaged.

We focused on the genes with positive fitness scores, as the overexpression of a gene that is important for a given process is usually expected to lead to a fitness advantage^17,46^, but we also examined the negative scores. To identify a subset of the effects that were likely to be reliable, we used three filters: the fitness effect was large relative to the variation between start samples (|score| >= 2); the fragments containing the gene showed consistent fitness across replicate experiments (using a *t* test); and the number of reads for those fragments was sufficient for the gene score to have little noise (see Methods). Effects that passed these filters were more likely to be consistent in replicate experiments (for example, see **Fig. 3d**). We considered an effect that passed these filters to be of high confidence if it was based on more than one fragment or if the gene had a large effect in another experiment for the compound. Overall, we identified 4,051 high-confidence effects, representing 813 of the 4,151 genes assayed (**Supplementary Table 3**). 400 different genes had a high-confidence fitness benefit when overexpressed in at least one condition, while the overexpression of 571 different genes led to a decrease in fitness in at least one condition. Nearly all experiments (153 of 155) had at least one gene with a high-confidence effect. By shuffling the measurements for each fragment in each experiment, we estimated a false discovery rate of less than 2% (Methods). Among the *E. coli* genes essential for viability when deleted^5^, 46 have a high-confidence benefit in at least in one experiment, demonstrating that gain-of-function approaches like Dub-seq can identify conditional phenotypes for genes that are not typically interrogated by loss-of-function approaches such as Tn-seq.

Some genes had positive fitness benefits across many conditions. In particular, five genes (*recA, galE, dgt, rcnA, fabB*) had high-confidence benefits in 10 or more different conditions. The most frequent benefits were found for *recA* and *galE*, which are disrupted in the DH10B derivative host strain we used^47^ (Methods). Even for pleiotropic genes, we find that they confer a more extreme beneficial phenotype in some conditions. For example, UDP-glucose 4-epimerase (*galE*) is highly beneficial to overexpress in the presence of 0.1 mM benzethonium chloride, with gene scores of +12 or +14 in two replicate experiments. All of *galE*’s other scores were under +5. Similarly, strand exchange and recombination gene *recA* shows high fitness scores of +6 in the presence of cisplatin, lomefloxacin and sodium chloride. In addition to these examples, we found that 32 genes provide growth advantage in 5 or more antibiotics, metals or other stress conditions, as compared to 241 genes showing growth benefit in just one condition (**Supplementary Table 3**).

Some of the Dub-seq experiments identified dozens of putatively beneficial genes. For example, with potassium acetate as the carbon source, we identified 56 genes that had high-confidence benefits in both of two replicate experiments (**Supplementary Table 3**). The two highest-scoring genes encode isozymes of aconitase (*acnA* and *acnB)*, which are part of the tricarboxylic acid cycle for oxidizing acetate^48^. But the relationship between the other beneficial genes and acetate catabolism is not obvious. As another example, in copper (II) chloride stress at 2 mM, 120 genes had high-confidence benefits. The genes with the highest scores were *envZ, mltD, citB/dpiA, mepM, mepS, cutC*, and other high-scoring genes encode outer membrane porins (*ompX, ompC, ompF*) or lipoprotein *nlpE* (**Supplementary Table 3**). Overexpression of most these genes is known to activate the complex regulatory network of envelope stress response via *cpxAR* and sigma-E^49,50^. Specifically, it is known that the copper tolerance phenotype observed in the case of *nlpE* overexpression is due to activation of Cpx pathway^51^. In the case of *cutC* overexpression, sigma-E driven small RNA *micL* encoded within *cutC* is overproduced, leads to targeted downregulation of *lpp* and sufficient for copper tolerance phenotype^52^. Finally, dozens of genes show growth benefits in the presence of the membrane-disrupting cationic surfactants benzethonium and benzalkonium. Most of these genes are involved in membrane lipid homeostasis, envelope stress response pathways and drug efflux systems (**Fig. 4, Supplementary Table 3**).

In total, we identified 41 instances where the Dub-seq fitness data is consistent with the known growth benefit imparted by the gene (**Supplementary Table 4**). These high confidence, known hits include genes encoding diverse functions such as efflux pumps, transporters, and regulators, as well as biosynthetic enzymes and small RNAs, each yielding enhanced fitness via diverse mechanisms. For example, overexpression of *cysE* (which encodes serine acetyltransferase) probably increases nickel tolerance 53 through increased glutathione biosynthesis^53^, while overexpression of *rnc* (which encodes RNase III) yields a growth benefit in nickel and cobalt stress, as it down-regulates the expression of *corA*, which encodes a transporter that mediates the influx of nickel and cobalt ions into the cell^54^.

In addition to the known cases, we also identified hundreds of genes that had not been previously associated with a tolerance phenotype in a specific condition, including *pssA, dcrA/sdaC, dcrB* in sisomicin; *pmrD* in aluminum; *treA, treB* and *phnM* in phosphomycin; sRNAs *chiX* in nickel and *ryhB* in zinc; and many genes of unknown function (**Fig. 4, Supplementary Table 3**). To follow up some of the novel observations, we assayed the growth of strains overexpressing the genes individually with and without added stress. We used *murA* overexpression as a test case, as this is known to confer resistance to phosphomycin^55^ (**Supplementary Fig. 5**). Growth curves confirmed that the overexpression of either *pssA* or *dcrB* confers resistance to the aminoglycoside antibiotic sisomicin, although the mechanism(s) by which this resistance is conferred remains unclear. The gene *pssA* encodes an essential phosphatidylserine synthase, while *dcrB* is a periplasmic protein with a role in phage infection^48^. Growth curves also confirm that the overexpression of the outer membrane protein MipA confers strong resistance to benzethonium chloride (**Supplementary Fig. 5**). *mipA* has previously been implicated in the resistance to other antibiotics^56^.

Gene overexpression can also decrease host fitness^16,17,46^ and may indicate important function for those gene products. We identified 570 genes with a high-confidence negative effect on fitness in at least one experiment (**Supplementary Table 3**). Some of these genes appear to be more generally toxic when overexpressed or have a global regulatory role and compromise host fitness in multiple conditions. 24 genes had detrimental effects on fitness in 10 or more different conditions (*ampH, arcZ, aroK, crr, gadY, hfq, hha, htpX, hupB, iraP, metJ, mtlA, nupG, rpoS, ruvA, tsx, wecA, ybjT, yceG, ydgA, ydjN, yibN, yjdC*, and *zinT*). Conversely, some genes have negative gene scores in only one or a handful of conditions. For example, consistent with earlier studies we found that overexpression of *glpT* or *uhpT* increases susceptibility to phosphomycin^57^. These results also agree with clinical data, which shows that the main cause of phosphomycin resistance in patients is the down-regulation of GlpT via down-regulation of cAMP.^57^ Accordingly, we also found that overexpression of *cpdA* (which encodes an enzyme that hydrolyzes cAMP) enhances fitness under phosphomycin stress (**Fig. 4**).

Finally, we analyzed our data for ‘epistatic’ instances where multiple genes on a fragment are necessary for the observed phenotype. Specifically, we searched for evidence of synergy between genes by analyzing scores for fragments containing more than one gene that are significantly greater than the inferred sum of score of the constituent genes (Methods). In total, we found 6 high scoring epistatic-effect cases across 52 conditions in our Dub-seq dataset (*fetA-fetB* on nickel, *ampD-ampE* on benzethonium, *ackA-pta* on D-lactate, *arcA-yjjY* on sisomicin, *hns-tdk* on phosphomycin and *yfiF-trxC* on potassium acetate (**Supplementary Fig.6abc**)). Among these, 3 gene-pairs have related functions (*fetA-fetB* form a complex, *pta-ackA* encode enzymes that catalyze adjacent reactions in the catabolism of lactate, and *ampD-ampE* are thought to be a signaling pathway^48^) and our data indicates, together they provide a larger growth benefit. Specifically, overexpression of *fetAB* together has been shown to improve survival during nickel stress^58^.

### Comparison to loss-of-function fitness data

Integrating large-scale genetic gain and loss of function can provide added specificity to biological insights. For instance, genes with resistance phenotypes when overexpressed and sensitivity phenotypes when deleted are often specifically involved in the condition of interest, as demonstrated by studies identifying drug targets in yeast^59^ or identifying small RNA regulators^60^ or antibiotic resistance factors in bacteria^61^. Furthermore, genes with opposing loss and gain-of-function phenotypes for stress compounds are more likely to be true resistance determinants as opposed to genes that have indirect effects when overexpressed^16^. For 45 of the conditions that we profiled in this study with Dub-seq, we can systematically compare these phenotypic consequences of overexpression to loss-of-function mutations as determined by random barcode transposon site mutagenesis^15^. The two data sets studied the same growth media and compounds, but not necessarily at the same concentrations, and they used different strains of *E. coli* (DH10B or BW25113). Across these 45 conditions, we identified 625 high-confidence benefits of overexpression (or 0.3% of gene-condition pairs). Of the 625 high-confidence benefits, 480 are for genes with RB-TnSeq data, and in 62 cases (12%), that loss of function led to a significant disadvantage (RB-TnSeq fitness < −1 and *t* < −4, where *t* is a t-like test statistic^13^). By chance, we would expect just 2.5% agreement, which is significantly less (*P* < 10^-15^, chi-squared test of proportions). Overall, we found moderate overlap between genes that are beneficial when overexpressed and important for fitness when disrupted (**Supplementary Table 3**).

To illustrate the biological insights that can be derived by systematically comparing gain and loss-of-function data on a genomic scale, we present 3 examples: growth in the presence of elevated nickel, cobalt, or sodium chloride (**Fig. 5abc**). Under each condition, we find that a number of genes that are both necessary for resisting the stress when knocked-out and sufficient for a resistance phenotype when singly overexpressed. These instances include known examples such as the aforementioned metal exporter RcnA^45^ and RNase III for cobalt and nickel tolerance^54^, as well as the osmolyte transporter ProP^62^ and envelope biogenesis factor YcbC (ElyC)^63^ for tolerance to osmotic stress imposed by sodium chloride. (In our Dub-seq data, *proP* and *ycbC* failed to pass the filters for high-confidence effects). In addition to these known examples, there are more novel observations (**Fig. 5abc**). Under nickel and cobalt stress, the uncharacterized protein YfgG (DUF2633) is important for tolerance, a finding that is supported by RB-Tnseq data^15^ and by individual growth curve analysis of an *yfgG* overexpression strain (**Fig. 5d**). While the precise biochemical function of YfgG is unclear, a close homolog of this protein in *Klebsiella michiganensis* is also important for fitness under nickel and cobalt stress^15^. As a second example, we find that ProY is important for nickel resistance. A ProY homolog in the related bacterium *K. michiganensis* is also important for nickel resistance^15^. Using individual strain growth curve analysis, we confirmed that overexpression of *proY* alone can confer nickel resistance to *E. coli* (**Fig. 5e**). While ProY is currently annotated as a cryptic proline transporter, we suspect that its function is to transport histidine as it can suppress histidine auxotrophy^25^ and homologs of this protein are required for histidine utilization in other bacteria^15^. In light of this, we speculate that the nickel resistance phenotype of ProY is due to increased sequestration of nickel ions by a higher intracellular concentration of histidine. As a final example, we found that the porphyrogen oxidase YfeX confers sodium chloride resistance in *E. coli*, a finding confirmed by an individual growth curve analysis (**Fig. 5f**). While we are unsure how this protein manifests this phenotype, we note that yfeX homologs are important for resisting sodium chloride in multiple bacteria^15^. We have provided a general working hypothesis for many of other genes with high fitness scores in **Supplementary Table 5.**

**Figure 5.**
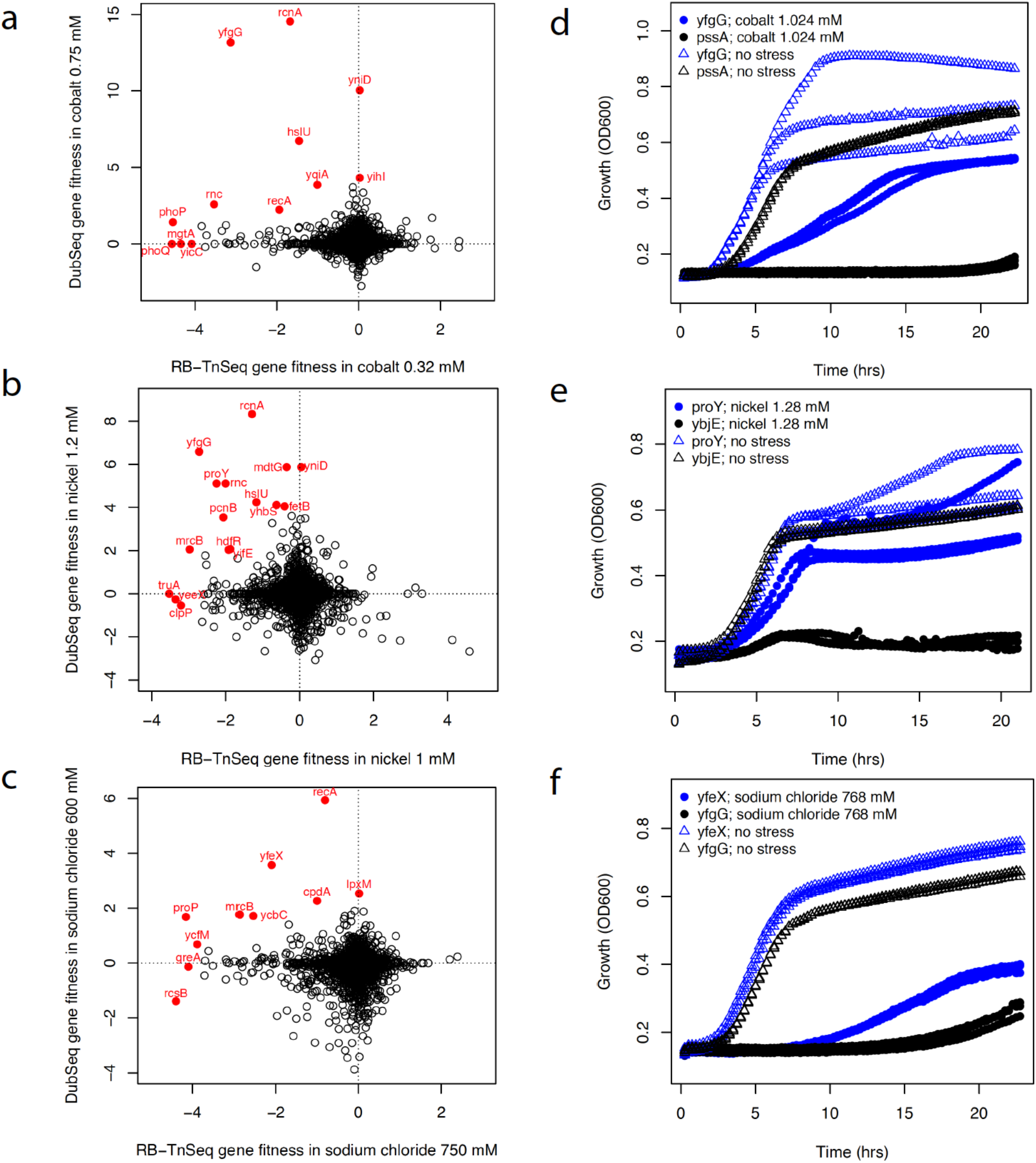
Comparing genome-wide loss and gain-of-function phenotype data. Comparison of RB-TnSeq fitness data^15^ (x-axis) and Dub-seq gene fitness data for *E. coli* genes under growth with inhibitory concentrations of cobalt (a), nickel (b), and sodium chloride (c). Selected genes are highlighted. (d) Growth of *E. coli* overexpressing *yfgG* under cobalt stress; *pssA* is a control. (e) Growth of *E. coli* overexpressing *proY* under nickel stress; *ybjE* is a control. (f) Growth of *E. coli* overexpressing *yfeX* under sodium chloride stress; *yfgG* is used as a control.

## DISCUSSION

Here we describe Dub-seq, a technology for performing parallelized gain-of-function fitness assays across diverse conditions. Dub-seq couples shotgun cloning of random DNA fragments with competitive fitness assays to assess the phenotypic importance of the genes contained on those fragments in a single tube assay. We demonstrate that Dub-seq is reproducible, economical, scalable, and identifies both known and novel gain-of-function phenotypes. By decoupling the library creation and characterization step from the screening step with BarSeq, Dub-seq provides a quantitative and rapid tool for experimentally assessing gene function via overexpression phenotypes of DNA cloned into an expression vector. This approach can improve overall repeatability and reproducibility of genome-wide gain-of-function experiments, and facilitate open distribution of libraries among researchers^64^.

In this proof-of-concept study, we generated a Dub-seq library of *E. coli* genomic DNA in a broad-range expression vector and assayed the phenotypic importance of overexpressing cloned genes using *E. coli* as the host bacterium. From 152 genome-wide assays, we identified 400 different genes with a high-confidence fitness benefit when overexpressed in at least one experimental condition. The majority of these gene-phenotype associations have not previously been reported including, as far as we know, for *yfgG, proY*, and *yfeX* (**Supplementary Table 3**). We found 241 genes confer a fitness benefit in just one condition, indicating a condition-specific phenotype. Overall, 32 genes enhanced fitness in 5 or more conditions, suggesting their broader role in host fitness and importance in cross-resistance phenotypes observed between metals, antibiotics, antiseptics and other stresses^65^. Dub-seq recapitulated 41 known instances of positive fitness effects, wherein the fitness phenotypes stem from diverse mechanisms, including overexpression of a compound target, active efflux of heavy metals, decreased uptake of metals and antibiotics, increased uptake of nutrients, and the regulatory effects of both protein-coding genes and small RNAs. We also identified enhanced susceptibility due to overexpression. Finally, we show that systematically comparing gain and loss-of-function datasets provide additional insights into those genes that are both necessary and sufficient for stress tolerance phenotypes.

Dub-seq can be readily extended to DNA from other sources and many cultured bacteria could be adapted as hosts for the genome-wide fitness assays. In particular, our vectors should be suitable to build Dub-seq libraries of microbial isolates and can be mobilized to new bacteria via conjugation because of its broad-host range replication origin. By using other hosts, we can overcome gene expression and toxicity issues associated with expressing heterologous DNA in model hosts^34-36^. To extend the Dub-seq methodology for functional profiling of DNA isolated from the environment, we would need to generate a higher diversity of barcoded vectors so that we would have a large library of unique barcode pairs and the largest percentage of metagenomic diversity can be captured and mapped confidently. In addition, to ensure reliable expression of heterologous genes, a number of approaches can be used to activate transcription or translation of genes encoded within foreign DNA^34,42,66^.

In this work, we generated a Dub-seq library with a ~2.6 kb insert size and therefore by design, the library only covers fragments encoding 2-3 genes on an average. Therefore, phenotypes that are only conferred by the activity of a larger group of genes (such as multisubunit complexes) will not be detected. Nevertheless, we did detect 6 instances of ‘epistatic’ interactions in which two neighboring genes show greater fitness score as gene-pairs than the inferred sum of score of the individual genes. By adapting the Dub-seq strategy to fosmids, cosmids and bacterial-artificial-chromosomes, future efforts can clone larger size genomic fragments to create Dub-seq libraries for the discovery of activities encoded by multiple genes, including secondary metabolites.

Given the increasing knowledge gap between genomic sequence and function, and the limited ability of computational approaches to accurately predict gene function from sequence, high-throughput experimental methods are needed to assign gene function and resolve roles of uncharacterized genes. Recently, a number of loss-of-function methods have been developed^5-8,10-14^, but only a fraction of genes from genetically tractable microbes can be readily annotated with a specific function using these approaches. We envision that multiple, complementary experimental approaches that can be applied *en masse* are ultimately necessary to uncover the roles of most poorly annotated genes from microbial isolates and microbiomes. The Dub-seq approach we presented here is another valuable tool in this toolkit.

## Author contributions

V.K.M., A.M.D. and A.P.A. conceived the project. V.K.M., A.M.D., A.P.A., supervised the project. V.K.M. led the experimental work. P.S.N. led the computational work. V.K.M., A.M.D., T.K.O., M.C. and S.C. collected data. V.K.M., P.S.N., M.N.P. and A.M.D. analyzed the fitness data. M.N.P. and A.P.A. provided advice on data processing and modeling. V.K.M., P.S.N., M.N.P., A.M.D. and A.P.A. wrote the paper.

## Acknowledgements

We thank Mahek Modi and Aaron Gupta for assisting in the initial stage of this project. The initial concepts for this project were developed by Biodesign project supported by the Office of Science (BER), U.S. Department of Energy, DE-SCOOO8812. The implementation was funded by ENIGMA, a Scientific Focus Area Program at Lawrence Berkeley National Laboratory, supported by the U.S. Department of Energy, Office of Science, Office of Biological and Environmental Research under contract DE-AC02- 05CH11231.

## METHODS

### Strains and growth conditions

*Escherichia coli* BW25113 was purchased from the *E. coli* Genetic Stock Center. All plasmid manipulations were performed using standard molecular biology techniques^67^. All enzymes were obtained from New England Biolabs (NEB) and oligonucleotides were received from Integrated DNA Technologies (IDT). *Escherichia coli* strain DH10B (DH10B derivative, NEB 10-Beta) was used for plasmid construction and as host for Dub-seq fitness assays. Unless noted, all strains were grown in LB supplemented with 30 μg/ml chloramphenicol at 37°C and shaking at 200 rpm. The primers, plasmids and strains used in this study are listed in **Supplementary Tables 6, 7** and **8** respectively.

### Construction of dual barcoded Dub-seq vector

To construct a double barcoded vector, we used pFAB5477 an in-house plasmid with 68 pBBR1 replication origin and a chloramphenicol resistance marker^68^. pBBR1 based broad-host plasmids are relatively small, mobilizable and have been widely used for a variety of genetic engineering applications in diverse microbes^69^. To insert a pair of DNA barcodes on the plasmid we used phosphorylated oFAB2853 and oFAB2854 primers to amplify the entire plasmid pFAB5477, removed the plasmid backbone using DpnI (as per manufacturing instructions, NEB), and ligated the amplified and pure product using T4 ligase (as per manufacturing instructions, NEB). The random N’s in oFAB2853 and oFAB2854 (**Supplementary Table 6**) represent the UP and DOWN barcode sequences. The ligated product, pFAB5491, was column purified using the Qiagen PCR purification kit, transformed into DH10B electro-competent cells (NEB 10-Beta *E. coli* cells, as per manufacturing instructions, NEB) and transformants were selected on LB-agar plates supplemented with 30 ug/ml chloramphenicol. The next day, ~250,000 colony forming units (CFU) were estimated and scraped together into 20 ml LB with 30 ug/ml chloramphenicol. The culture library was diluted to an optical density at 600 nm (OD600) of 0.2 in fresh LB medium supplemented with 30 ug/ml chloramphenicol and grown to a final OD600 of ~1.2. We added glycerol to a final concentration of 15%, made multiple 1 ml glycerol stocks, and stored them at −80°C. We also collected cell pellets to prepare plasmid DNA of pFAB5491 for further characterization of the library (BPseq).

### BPseq to characterize dual barcoded Dub-seq vector

To associate the pair of DNA barcodes, we performed Barcode-Pair sequencing (BPseq) of the plasmid pFAB5491 library. For deep coverage of the library, we performed 10 different PCR reactions using primers VM_barseq_P1 and VM_Barseq-P2. The forward primers VM_Barseq-P2 contains different 6-bp TruSeq indexes, and were automatically demultiplexed by the Illumina software.

We performed PCR in a 100-ul total volume with 5 ul common reverse primer VM_barseq_P1 (4 uM), 5 ul forward primer VM_Barseq-P2 _IT001 to IT010 (4 uM), 38 ul of sterile water, 2 ul template pFAB5491, and 50 ul of 2X stock of Q5 DNA Polymerase mix (500 ul of 2X stock of Q5 DNA Polymerase mix consists of 200 ul Q5 buffer, 20 ul dNTP, 50 ul DMSO, 10 ul Q5 DNA Polymerase enzyme and 220 ul water) under following PCR conditions: 98°C for 4 minutes, followed by 15 cycles of 30 sec at 98°C, 30 sec at 55°C, 30 sec at 72°C and final extension at 72°C for 5 minutes. Finally, we ran the PCR products on an analytical gel to confirm amplification. We pooled equal volumes (10 ul) of BarSeq PCR products, purified the combined product using Qiagen PCR purification kit, and eluted in 40 ul of sterile water. We quantified the DNA product with a Qubit double-stranded DNA (dsDNA) high-sensitivity (HS) assay kit (Invitrogen). The BPseq samples were sequenced first on Illumina MiSeq and then HiSeq 2500: both with 150 bp single-end runs.

### BPseq data analysis

BPseq reads were analyzed with *bpseq* script from the *Dub-seq* python library with default parameters (code available at https://github.com/psnovichkov/DubSeq). The script looks for the common flanking sequences around each barcode (UP and DOWN) and requires an exact match of 9 nucleotides on both sides. By default, these flanking sequences may be up to 2 nucleotides away from their expected positions. The script also requires that each position in each barcode have a quality score of at least 20 (that is, an estimated error rate of under 1%). This gives an initial list of pairs of barcodes with the correct length and reliable sequence quality.

We applied two additional filters to minimize the number of erroneous barcode pairs that can be caused by PCR artifacts or sequencing errors. First, we check whether a given barcode can be a result of a single nucleotide substitution introduced in a real barcode and filter out all such barcodes. We perform a pairwise sequence comparison of all extracted barcodes (UP and DOWN barcodes are treated separately) and search for “*similar* barcodes. Two barcodes are considered to be *similar* if they are different by only one nucleotide. A given barcode passes the filter if it does not have similar barcodes or it is at least two times more frequent than the most abundant similar barcode.

Second, we check whether a given barcode pair can be a result of chimeric PCR and filter out all such pairs. As the region between and around UP and DOWN barcodes are identical in all plasmids in our library, we expected artifacts from formation of chimeric BPseq PCR products^13^. We perform a pairwise comparison of all barcode pairs and search for “*related”* pairs. Two barcode pairs are considered to be related if they have either the same UP or DOWN barcodes. The presence of the same UP (or DOWN) barcode in multiple barcode pairs is potentially a sign of chimeric PCR. To distinguish the true barcode pair from the chimeric one, we check the frequency of all the related barcode pairs. A given barcode pair passes the filter and is considered to be non-chimeric if it does not have related pairs or it is at least two times more frequent than the most abundant related barcode pair. As a result, the ‘reference set’ of barcode pairs is created. From the BPseq step we obtained 5,436,798 total reads. Among these, total usable reads (reads that support barcode pairs from the reference set) were 2,933,702 and represent about 54% of total reads.

### Dub-seq vector preparation for cloning genomic fragments

To prepare the Dub-seq vector pFAB5491 for cloning, we made 900 ul or about 100 ug of plasmid preparation (Qiagen plasmid miniprep kit), and performed two rounds of PmiI digestion. Restriction digestion reaction included 900 ul (total 100 ug) of pFAB5491 plasmid, 100 ul PmiI enzyme, 400 ul 10X cutsmart buffer, and water to make up the volume of 4000 ul. We incubated the reaction at 37°C on a heating block for 4 hours and then checked the reaction progress on an analytical 1% agarose gel. To dephosphorylate the restriction-digested vector, we added 1 unit of rSAP for every 1 pmol of DNA ends (about 1 μg of a 3 kb plasmid), and incubated at 37°C for 2 hours in a PCR machine. We stopped the reaction by heat-inactivation of rSAP and restriction enzyme at 70°C for 20 minutes. The cut and dephosphorylated vector library was then gel purified (Qiagen gel extraction kit). To remove any uncut vector, we repeated the entire process of restriction digestion, dephosphorylation, and purification. The final concentration of cut and pure barcoded vector library used for cloning genome fragments was about ~30 ng/ul.

### Construction of *E. coli* Dub-seq library

To construct Dub-seq library of *E. coli* genomic fragments, we extracted *E. coli* BW25113 genomic DNA and 1 ug was fragmented by ultrasonication to an average size of 3000 bp with a Covaris S220 focused ultrasonicator. The sheared genomic DNA was then gel purified and end-repaired using End-IT kit (Epicentre, as per manufacturer instruction). Briefly the 50 ul reaction included: 34 ul sheared DNA (1.0 ug total), 5 ul ATP 10 mM, 5 ul dNTP mix (10 mM), 5 ul EndIt buffer 10X and 1-2 ul EndIT enzyme. We incubated the reaction at room temperature for 45 mins, and inactivated the enzyme by incubating the reaction at 70°C for 10 minutes. The end-repaired genome fragments were purified with PCR clean-up kit (Qiagen), and quantified on Nanodrop.

The end-repaired genomic fragments were then ligated to the restriction-digested, sequence-characterized dual barcoded backbone vector (pFAB5491) at 8:1 insert:vector ratio using Fast-link Ligase enzyme (Epicentre, as per manufacturer instruction). The total 60 ul ligation reaction consists of 4 ul of restriction-digested pFAB5491, 20 ul End-repaired DNA, 3 ul ATP (10 mM), 6 ul 10X ligase buffer, 19 ul water and 8 ul Fast-link-ligase. The ligation was incubated overnight (18 hrs) at 16°C, inactivated at 75°C for 15 minutes, and purified using PCR purification kit (Qiagen).

For transforming the ligation reaction, 60 ul of column-purified ligation reaction was mixed gently with 1500 ul of NEB DH10B electrocompetent cells on ice and then the mix was dispensed 60 ul per cuvette. Electroporation was done using parameters supplied by NEB. Transformed cells were recovered by adding 1 ml SOC recovery media (as per competent cell manufacturer instruction, NEB). We pooled all recoveries and added additional 10 ml of fresh SOC. Transformants were then incubated at 37°C with shaking for 90 minutes. We spun down the pellets and resuspended the pellet in 6 ml SOC. Different volumes of 6 ml resuspended pellets were then plated on overnight-dried bioassay plates (Thermo Scientific # 240835) of LB agar supplemented with 30 ug/ml chloramphenicol. We also did dilution series for estimating CFUs.

We determined the number of colonies required for 99% coverage of *E. coli* genome using the formula N = ln(1 −0.99)/ln(1-(Insert size/Genome Size)) to ensure that genome fragments are present in the cloned library^70^. For example, to cover the *E. coli* genome (of size 4.7 Mb) with fragments of 3 kb, we need about 4,610 strains for 99% coverage. We collected ~40,000 colonies by scraping the colonies using a sterile spatula into 20 ml LB supplemented with 30 ug/ml chloramphenicol in a 50 ml Falcon tube and mixed well. This *E. coli* Dub-seq library was then diluted to an optical density at 600 nm (OD600) of 0.2 in fresh LB supplemented with 30 ug/ml chloramphenicol and grown to a final OD600 of ~1.2 at 37°C. We added glycerol to a final concentration of 15%, made multiple stocks of 1 ml volume, and stored the aliquots at −80C. We also made cell pellets to store at −80°C and to make large plasmid preparation (Qiagen) for BAGseq library preparation.

### BAGseq to characterize barcoded genomic fragment junctions

We characterized the final plasmid library pFAB5516 using a TnSeq-like protocol^13^, which we call Barcode-Association-with Genome fragment sequencing or BAGseq. BAGseq identifies the cloned genome fragment and its pairings with neighboring dual barcodes. This step of associating the dual barcodes with each library of genomic fragments is only done once (by deep sequencing) and used as a reference table to derive connections between observed functional/fitness traits with specific cloned genomic fragment (Fig. 1).

To generate Illumina-compatible sequencing libraries to link both UP and DOWN random DNA barcodes to the ends of the cloned genome fragments, we processed two samples per library. The plasmid library (1 ug) samples were fragmented by ultrasonication to an average size of 300 bp with a Covaris S220 focused ultrasonicator. To remove DNA fragments of unwanted size, we performed a double size selection using AMPure XP beads (Beckman Coulter) according to the manufacturer’s instructions. The final fragmented and size-selected plasmid DNA was quality assessed with a DNA 1000 chip on an Agilent Bioanalyzer. Illumina library preparation involves a cascade of enzymatic reactions, each followed by a cleanup step with AMPure XP beads. Fragmentation generates plasmid DNA library with a mixture of blunt ends and 5’ and 3’ overhangs. End repair, A-tailing, and adapter ligation reactions were performed on the fragmented DNA using the NEBNext DNA Library preparation kit for Illumina (New England Biolabs), according to the manufacturer’s recommended protocols. For the adapter ligation, we used 0.5 ul of a 15uM double-stranded Y adapter, prepared by annealing Mod2_TS_Univ (ACGCTCTTCCGATC*T) and Mod2_TruSeq (Phos-GATCGGAAGAGCACACGTCTGAACTCCAGTCA). In the preceding oligonucleotides, the asterisk and Phos represent phosphorothioate and 5’ phosphate modifications, respectively.

To specifically amplify UP barcodes and neighboring genomic fragment terminus by PCR, we used the UP-tag-specific primer oFAB2923_Nspacer_barseq_universal, and P7_MOD_TS_index1 primer. For the DOWN-tag amplification we used oFAB2924_ Nspacer_barseq_universal and P7_MOD_TS_index2 primer. For the BAGseq UP barcode and DOWN barcode site enriching PCR, we used JumpStart Taq DNA polymerase (Sigma) in a 100 ul total volume with the following PCR program: 94°C for 2 minutes and 25 cycles of 94°C 30 seconds, 65°C for 20 seconds, and 72°C for 30 seconds, followed by a final extension at 72°C for 10 minutes. The final PCR product was purified using AMPure XP beads according to the manufacturer’s instructions, eluted in 25 ul of water, and quantified on an Agilent Bioanalyzer with a DNA-1000 chip. Each BAGseq library was then sequenced on the HiSeq 2500 system (Illumina) with a 150 SE run to map UP and DOWN barcodes to genomic inserts in the Dub-seq *E. coli* library.

### BAGseq data analysis

BAGSeq reads were analyzed with *bagseq* script from the *Dub-seq* python library with default parameters (code available at https://github.com/psnovichkov/DubSeq). Fastq files for UP and DOWN barcodes with associated (cloned) genomic fragments are processed separately. For each read, the script looks for the flanking sequences around a barcode and requires an exact match of 9 nucleotides on both sides and a minimum quality score of 20 for each nucleotide in a barcode. The sequence downstream of the identified barcode is considered to be a candidate genomic fragment and is required to be at least 15 nucleotides long for further processing. As a result, the initial list of the extracted barcodes and candidate genomic fragments is constructed.

All extracted genomic fragments were compared to the *E. coli* genome sequence with BLAT using default parameters. Only hits with alignment block size of at least 15 nucleotides and at most one indel were considered. It is also required that the extracted genomic fragment is mapped to one location in the genome. Thus, mappings to repeat regions were ignored. We applied two additional filters to minimize the number of erroneous associations between barcode and genomic location. First, we applied the same type of filter that we use for the analysis of BPSeq reads to filter out barcodes with a 1-nucleotide error.

Second, the same barcode can be associated with different genomic fragments because of PCR artefacts (chimeras) or because multiple fragments were cloned between the same pair of barcodes. To filter out erroneous barcode mappings, the number of reads supporting different locations for the same barcode were calculated. To distinguish the true location from the false one, the frequency of the most abundant location (the number of supported reads) was compared with frequencies of all other locations for the same barcode. A given association between the barcode and the genomic location is considered to be true if the barcode does not have any other associated locations or the abundance of this association is at least two times more frequent than any other associations for the same barcode. As a result, the reference set of associations between UP (and separately for DOWN) barcodes and genomic locations is created, which we call ‘BAGseq reference set’.

The BPseq reference set of barcode pairs and BAGseq reference set are combined together to associate pairs of barcodes with genomic regions (to create the final ‘Dub-seq reference set’). This step is done using the *bpag* script from the *Dub-seq* python library with default parameters. For each BPseq barcode pair, the script checks if the associations between UP and DOWN barcodes with genomic locations are present in the BAGSeq reference set. If both UP and DOWN barcodes (from BPseq reference set) are mapped to the genome, then the script checks the length of the region between the mapped locations and requires it to be between 100 nt and 6 kb. As a result, the final Dub-seq reference list of barcode pairs associated with genomic regions is created. Among total 10,600,088 reads for UP barcodes, usable reads were 3,884,931 (BAGseq UP barcode reads supporting the Dub-seq reference set), representing about 36.65% of total reads, whereas for total 9,671,635 reads for DOWN barcodes, usable reads were 2,499,399, representing about 25.84% of total reads (BAGseq DOWN barcode reads supporting the Dub-seq reference set).

### Competitive growth experiments

For genome-wide competitive growth experiments, a single aliquot of the Dub-seq library in *E. coli* DH10B was thawed, inoculated into 25 ml of LB medium supplemented with chloramphenicol (30 ug/ml) and grown to mid-log phase. At mid-log phase, we collected cell pellets as a common reference for BarSeq (termed start or time-zero samples) and we used the remaining cells to set up competitive fitness assays under different experimental conditions at a starting OD600 of 0.02. For carbon source growth experiments, we used M9 defined medium supplemented with 0.3 mM L-leucine (as DH10B is auxotrophic for L-leucine)^47^ and chloramphenicol. For experiments with stress compounds, we used an inhibitory but sublethal concentration of each compound, as determined previously^15^. All stress experiments were done in LB with chloramphenicol. All pooled fitness experiments were performed in 24-well microplates with 1.2 mL of media per well and grown in a multitron shaker. We took OD readings periodically in a Tecan M1000 instrument to ensure that the cells were growing and to confirm growth inhibition for the stress experiments. The assayed Dub-seq library cell pellets were stored at −80C prior to plasmid DNA extraction.

### BarSeq

Plasmid DNA from Dub-seq library samples was extracted either individually using the Plasmid miniprep kit (Qiagen) or in 96-well format with a QIAprep 96 Turbo miniprep kit (Qiagen). Plasmid DNA was quantified with the Quant-iT dsDNA BR assay kit (Invitrogen). The BarSeq PCR of UP barcodes was done as previously described^13^ with ~50 ng of plasmid template per BarSeq PCR reaction. To quantify the reproducibility of both UP and DOWN barcodes in competitive growth experiments, we collected plasmid DNA from nickel and cobalt experiments, and amplified both UP and DOWN barcodes in two separate PCRs using the same plasmid library template. For BarSeq PCR of DOWN barcodes, we used universal-forward-primer DT_BarSeq_p1_FW and reverse primer DT_BarSeq_IT017. The PCR cycling conditions and purification steps were same as for the UP barcodes^13^. All experiments done on the same day and sequenced on the same lane are considered as a ‘set’.

### BarSeq data analysis and fragment score calculation

From HiSeq 4000 runs we obtained ~400 million of reads per lane, or 4.2 million reads per sample (for multiplexing 96 samples) typically >60% reads were informative after filtering out reads for sequencing errors and unmapped barcodes. BarSeq reads were analyzed with *barseq* script from the *Dub-seq* python library with default parameters. For each read, the script looks for the flanking sequences around each barcode and requires an exact match of 9 nucleotides on both sides and a minimum quality score of 20 for each nucleotide in a barcode. The number of reads supporting each barcode is calculated. We apply the same type of filter that we use for the analysis of BPSeq reads to filter out barcodes with single nucleotide substitutions relative to real barcodes (see BPSeq section). As a result, the list of barcode and their counts is created.

### Calculation of fragment scores (fScores)

Given a reference list of barcodes mapped to the genomic regions (BPSeq and BAGSeq), and their counts in each sample (BarSeq), we estimate fitness values of each genomic fragment (strain) using *fscore* script from the Dub-seq python library with default parameters. First, the script identifies a subset of barcodes mapped to the genomic regions that are well represented in the time-zero samples for a given experiment set. We require that a barcode have at least 10 reads in at least one time-zero sample to be considered a valid barcode for a given experiment set. Then the *fscore* script calculates fitness score only for the strains with valid barcodes.

Strain fitness (*f_i_*) is calculated as a normalized log_2_ ratio of counts between the treatment (condition or end) sample *s_i_* and sum of counts across all (start) time-zero *t_i_*

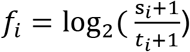

Then the strain fitness scores are normalized so that the median in each experiment is zero

### Calculating gene-score (gScore)

Given the fitness scores calculated for all Dub-seq fragments, we estimate a fitness score for each individual gene that is covered by at least one fragment. As mentioned in the Results, simply averaging the scores for the fragments that cover a gene gives spurious results for non-causative genes that are adjacent to a causative gene. To overcome this problem we modeled the fitness score of each fragment as the sum of the fitness scores of the genes that are completely covered by this fragment. Our model for estimating gene scores assumes that genes contribute independently to fitness, that most genes have little impact on fitness, and that intergenic regions have no effect on host fitness.

To estimate gene scores, we cannot use ordinary least squares (OLS), the most common type of regression, because of over fitting, which would produce unrealistic high positive and low negative scores for many genes. We also considered regularization methods (Ridge, LASSO, and ElasticNet), but these suffered from either too much shrinkage of fitness scores (biasing them towards zero) or failed to eliminate over fitting (see **Supplementary note**). Instead, we use Non-Negative Least Squares (NNLS) regression^71^, where the predicted gene scores are restricted to take only nonnegative values. If a gene with a potential benefit is next to (but not covered by) a fragment with negative fitness, most regression methods would inflate the benefit of the gene and assign a negative score to the nearby gene. NNLS instead ignores the (often noisy) negative scores for the nearby fragments. To estimate negative gene scores, we also used NNLS, but with the signs of the fragment scores flipped.

In our model, the expected fitness of a fragment is given by

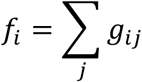

were *g_tj_* is a fitness score of a gene covered by *i*-th fragment completely. The NNLS minimizes

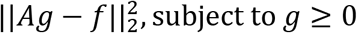

where *g* a vector of gene fitness scores to be estimated, *f* is vector of the “observed” fitness scores of fragments, *A* a matrix of ones and zeros defining which gene is covered by which fragment completely. Gene scores were calculated using the *gscore* script from the Dub-seq python library with default parameters, which uses the nnls function from the *optimize* package of the *scipy* python library.

### High-confidence gene scores and estimating the false discovery rate

We used several filters to identify gene scores that were likely to be of high-confidence and reliable. Whereas the non-negative regression was used to determine if the high fitness of the fragments covering the gene are due to this gene or a nearby gene, these filters were intended to ensure that the fragments covering the gene had a genuine benefit. The first filter was |gene score| >= 2, as such a large effect occurred just 4 times in 17 control comparisons between independently-processed but identical “start” samples (0.2 per experiment). In contrast the actual conditions gave 40 large effects per experiment on average (over 150 times more).

Second, we noticed that some genes had high scores because of a single fragment with a very high score. These fragments did not have high scores in replicate experiments, so their high scores might be due to secondary mutations. To filter out these cases, we performed a single-sample *t* test on the fragment scores (for the fragments that covered the gene) and required *P* < 0.05. This test asks if the mean is significantly different from a reference value. To handle uncertainty in the true centering of the fragment scores (which were normalized to have a median of zero), we considered the mean of all fragment scores for the experiment. We used this as the reference value (instead of zero) if this mean had the same sign as the gene’s score. This makes the filter slightly more stringent. If the gene has just one fragment, then we cannot apply the *t* test, so we instead require that |fragment score| be in the top 1% for this experiment.

Third, we checked that the effect was larger relative to the expected noise in the mean of the fragment scores that cover the gene. The expected noise for each fragment can be estimated as sqrt(1/(1+count_after) + 1/(1+count_start)) / ln(2). This approximation is derived from the best case that the noise in the counts follows a Poisson distribution. The expected noise for the mean of the fragment scores is then sqrt(sum(fragment_noise^2^)) / nfragments. Note that z = mean(fragment score) / noise would (ideally) follow the standard normal distribution. We use |z| >= 4 as a filter; with 4,303 genes being assayed, we would expect about 0.3 false positives per experiment.

“Filtered effects” (that passed all three filters) were considered to be reliable. Reliable effects were considered to be high-confidence if the gene was covered by multiple fragments. Because of the risk of secondary mutations, a measurement for a gene with a single fragment was only considered high-confidence if it was reliable and was also supported by a large effect (|score| >= 2) in another experiment for that compound.

The filtered effects were usually consistent across replicate experiments and represent reliable scores. We had two biological replicates for 64 of the 82 conditions (a compound at a given concentration) that we studied. Across these 64 pairs of replicate experiments, 85% of genes with filtered effects in one replicate were consistent (|score| >= 1.5 and the same sign) in the other replicate. Large effects (|score| >= 2) were more likely to replicate if they were filtered (85% vs. 59% otherwise). Among filtered effects for genes covered by more than one fragment, 39% of the effects that did not replicate were from a single condition (zinc sulfate stress at 1 mM). We did not identify any obvious issue for the data from this condition. In total, 4,303 genes are covered by at least one fragment, but there are only 4,151 genes with at least one gene score (adequate representation in at least one start sample).

To estimate the false discovery rate for high-confidence effects, we randomly shuffled the mapping of barcodes to fragments, recomputed the mean scores for each gene in each experiment, and identified high-confidence effects as for the genuine data. This shuffling test will probably overestimate the FDR because it assumes that all of the variability in the fragment scores is due to noise. Also, we used the mean score, rather than regression-based gene score, in this test. This might also lead to an overestimate of the FDR. We repeated the shuffle procedure 10 times. On average, each shuffled data set had 75 high-confidence effects, while the actual data had 4,051 high-confidence effects, so we estimated the false discovery rate as 75/4051 = 1.9%.

### Calculating gene-pair fitness score

Although our model assumes that the genes on a fragment contribute independently to fitness, there are cases where multiple nearby genes work together to confer a phenotype. For estimating such ‘epistatic’ synergistic fitness contribution by neighboring pair of genes, we included additional variables in our fitness calculation to account for the contribution of pairs of adjacent genes (and their intergenic regions). For a gene-pair to qualify to be valid hit, the score for the gene-pair has to be more than the individual gene scores from single-gene regression model, scores should be consistent across replicates and should be supported by more than one fragment. After manual filtering, we found 6 high scoring epistatic-effect instances where gene-pairs positively contribute to the host fitness under specific condition (**Supplementary Table 5**). Among these, 3 gene-pairs have related functions (*fetA-fetB* on nickel, *ampD-ampE* on benzethonium, *ackA-pta* on D-lactate^48^) and make biological sense. However, in the other 3 high scoring gene-pairs *arcA-yjjY, hns-tdk* and *yfiF-trxC*, each gene is divergently transcribed and the reason behind combined fitness phenotype is not obvious. We speculate, the fitness phenotype in these cases may be function of intergenic regions in addition to the encoded genes.

### Experimental validation of single genes

To experimentally validate some of top hits in our Dub-seq results we used the ASKA ORF collection^29^. The ASKA library consists of *E. coli* ORFs cloned on a pMB1 replication origin plasmid and driven by an IPTG-inducible promoter. We extracted individual ASKA ORF plasmids from the collection, sequence confirmed and transformed the plasmids into our assay strain *E. coli* DH10B. As the plasmid copy number and the strength of promoter and ribosome binding site used in the ASKA ORF collection is different from the broad-host pBBR1 plasmid system used in *E coli* Dub-seq library, we screened for an optimum IPTG levels to induce the expression of specific gene in order to study the host fitness. We grew the individual strains in 96-well microplates with 150 uL total volume per well. These plates were grown at 30°C with shaking in a Tecan microplate reader (either Sunrise or Infinite F200) with optical density readings every 15 minutes.

### Library visualization tools

We used the Dub-seq viewer tool from the *Dub-seq* python library (https://github.com/psnovichkov/DubSeq) to generate regions of the *E. coli* chromosome covering fragments (landscape mode) presented in **Fig 2a**. To generate fitness score plots as shown in **Fig. 3a and 3b**, and **Supplement Figs. 4, 6 and 7**, we used gene-browser mode. We used Circa software (OmGenomics) to generate genome coverage plot shown in **Fig. 2a**.

### Code and metadata availability

Code for processing and analyzing Dub-seq data is available at https://github.com/psnovichkov/DubSeq

Complete data from all experiments (read counts per barcode, fragment scores and gene scores) is deposited here: https://doi.org/10.6084/m9.figshare.6752753.v1

Link to website with supplementary information and bulk data downloads: href="http://morgannprice.org/dubseq18/

## SUPPLEMENTARY INFORMATION

### Supplementary Tables

**Supplementary Table 1.** List of 135 genes not represented in *E. coli* Dub-seq library

**Supplementary Table 2.** List of protein-coding genes with details on number of Dub-seq fragments covering the gene, and if the gene is essential (according to the Keio library^5^), has RB-TnSeq data^15^ and has Dub-seq data (this work).

**Supplementary Table 3.** Filtered gene scores for reliable effects in Dub-seq dataset and if they have representative data in RB-TnSeq mutant library^15^

**Supplementary Table 4.** List of genes whose high dosage is known to yield positive fitness effects

**Supplementary Table 5.** Novel gene-function associations with fitness score >=4; hypothesis and general notes

**Supplementary Table 6.** List of primers used in this work

**Supplementary Table 7.** List of plasmids used in this work

**Supplementary Table 8.** List of strains used in this work

Link to website with supplementary information: http://morgannprice.org/dubseq18/">

## Supplementary Figures

**Supplementary Fig. 1.**
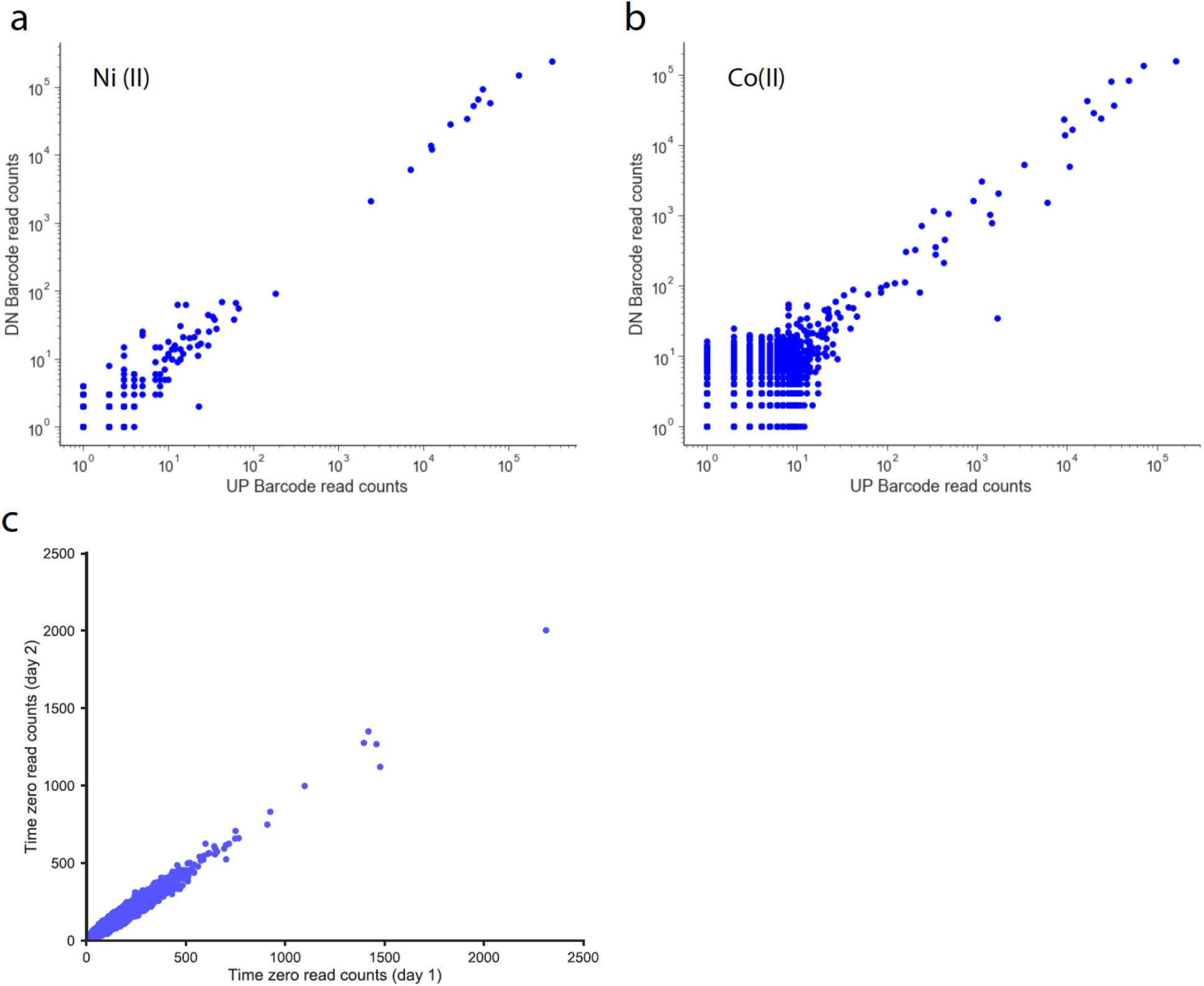
BarSeq reproducibility: Comparison of UP and DOWN barcode BarSeq reads for (a) Nickel and (b) Cobalt condition. (c) Comparison of UP barcode reads for two independent start (time-zero) samples.

**Supplementary Fig. 2.**
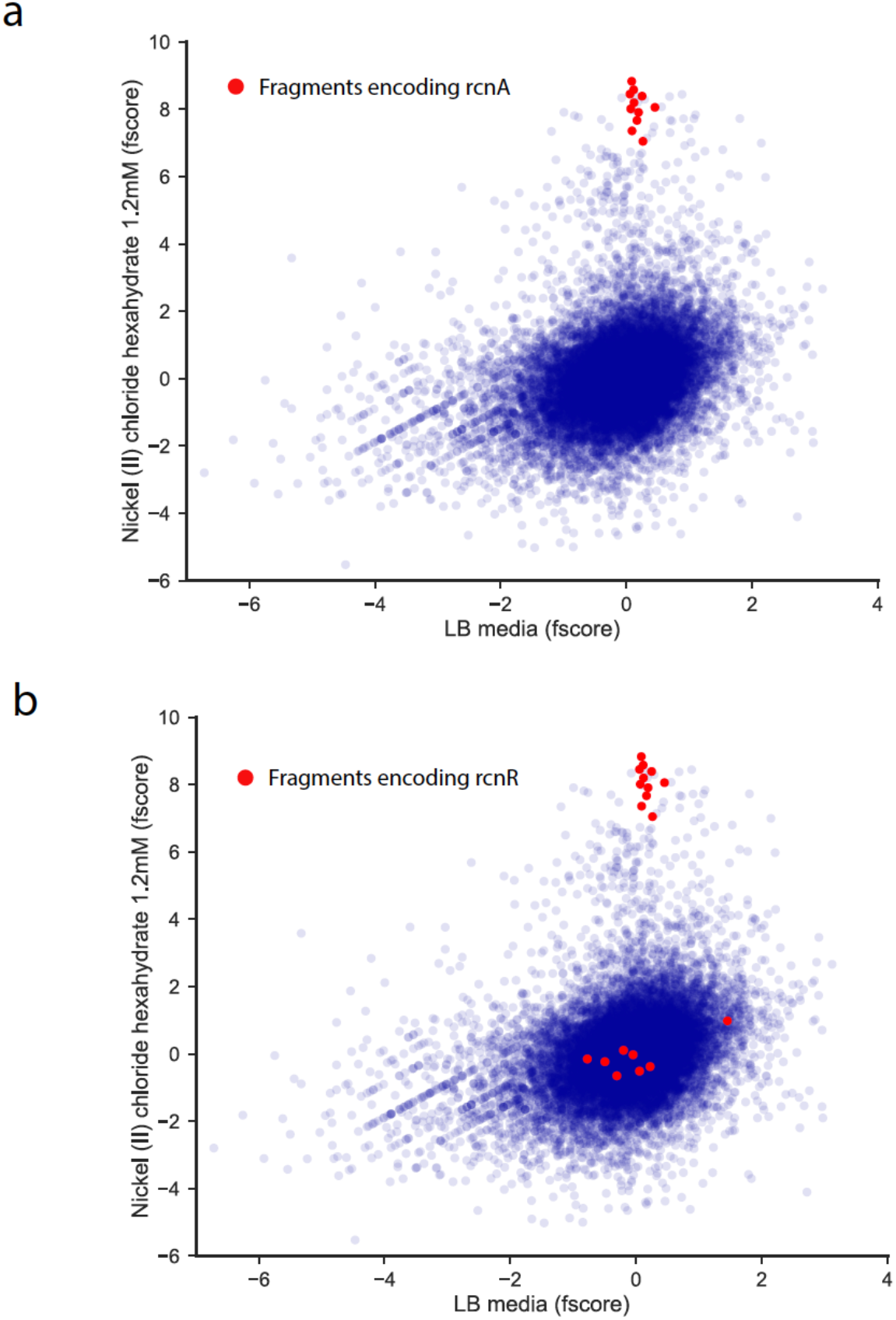
Fragment score comparisons: Fragment score (fscore) comparisons for all fragments in LB (x-axis) and LB with nickel (y-axis). (a) Fragments fully covering *rcnA* are highlighted in red. (b) Fragments fully covering *rcnR* are highlighted in red.

**Supplementary Fig. 3.**
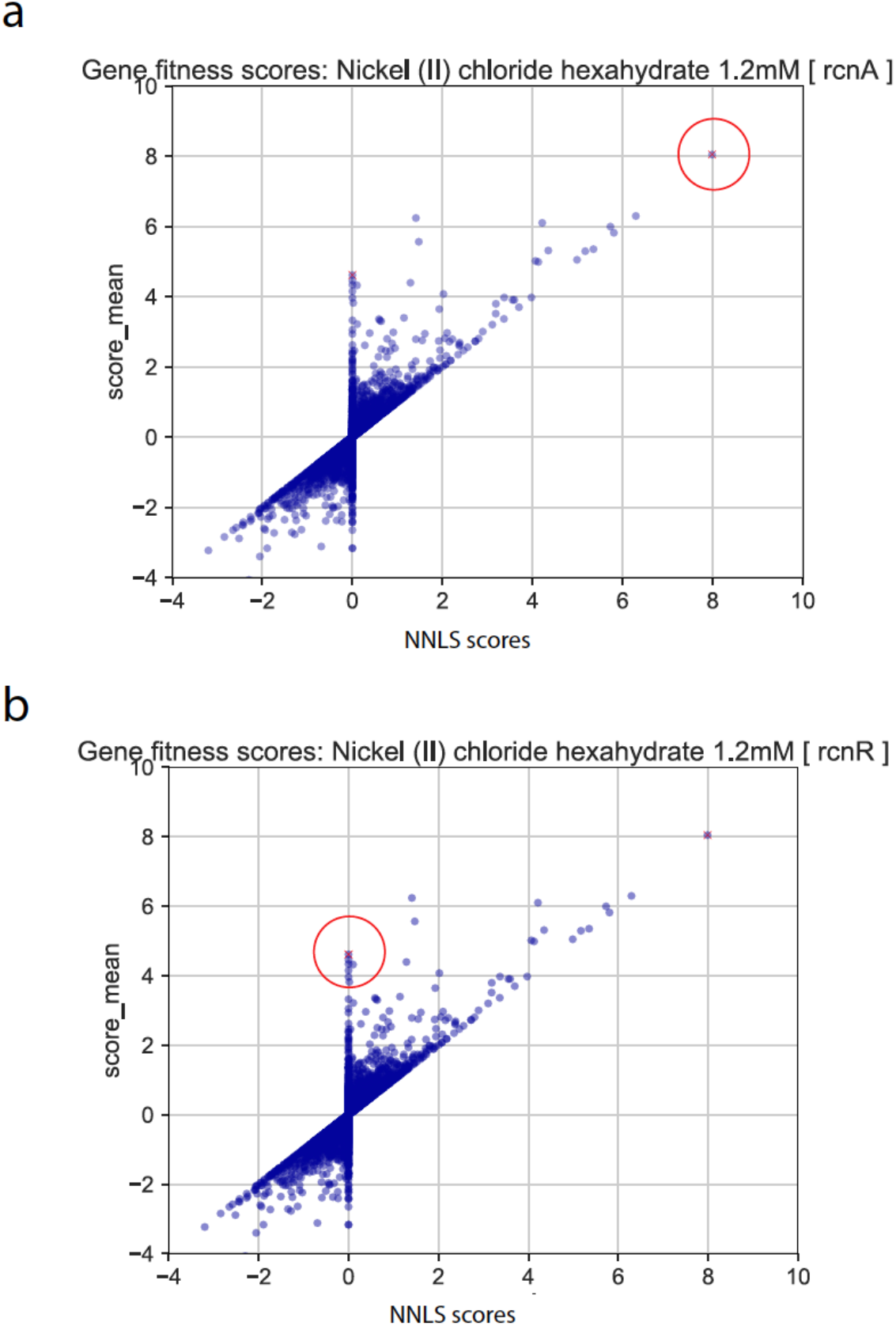
Comparison of gene scores from regression analysis and mean gene scores: Comparison between gene fitness scores calculated using Non-Negative Least Squares regression (NNLS) method and the mean score method under nickel stress (a) Fitness score for *rcnA* (red circle) (b) Fitness score for *rcnR* (red circle).

**Supplementary Fig. 4.**
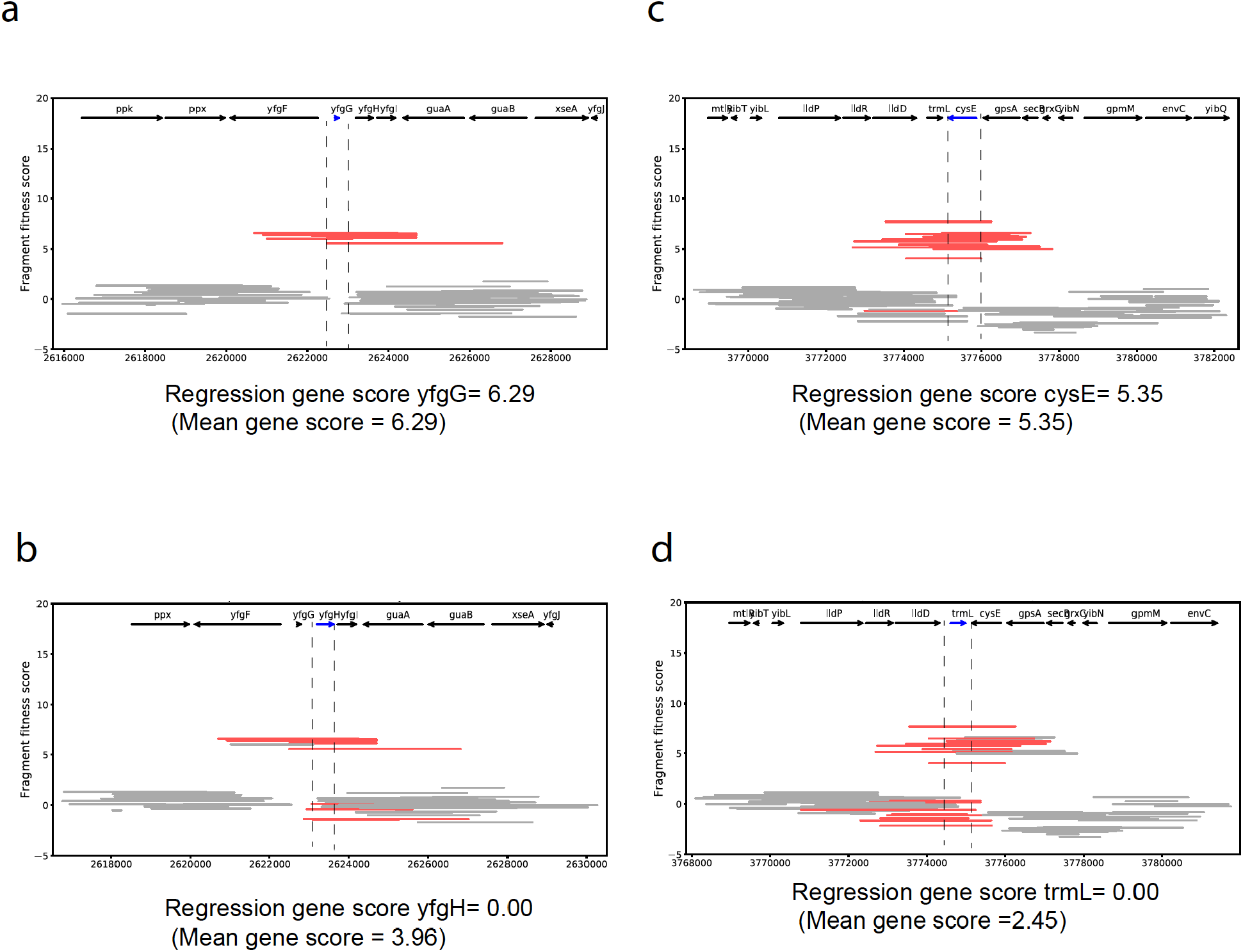
Fragment and gene Dub-seq scores: Dub-seq fragment (strain) data for different regions under elevated nickel stress (y-axis). Each line shows a Dub-seq fragment with those that completely cover the indicated gene are in red. The mean and regression scores for each indicated gene are shown below each plot. Compare scores for (a) *yfgG* with (b) *yfgH*, and (c) *cysE* with (d) *trmL.* Note that the mean and regression scores for *yfgH* and *trmL* are different. The mean score is incorrectly high for *yfgH* and *trmL* and is due to the presence of *yfgG and cysE* on a number of fragments.

**Supplementary Fig. 5.**
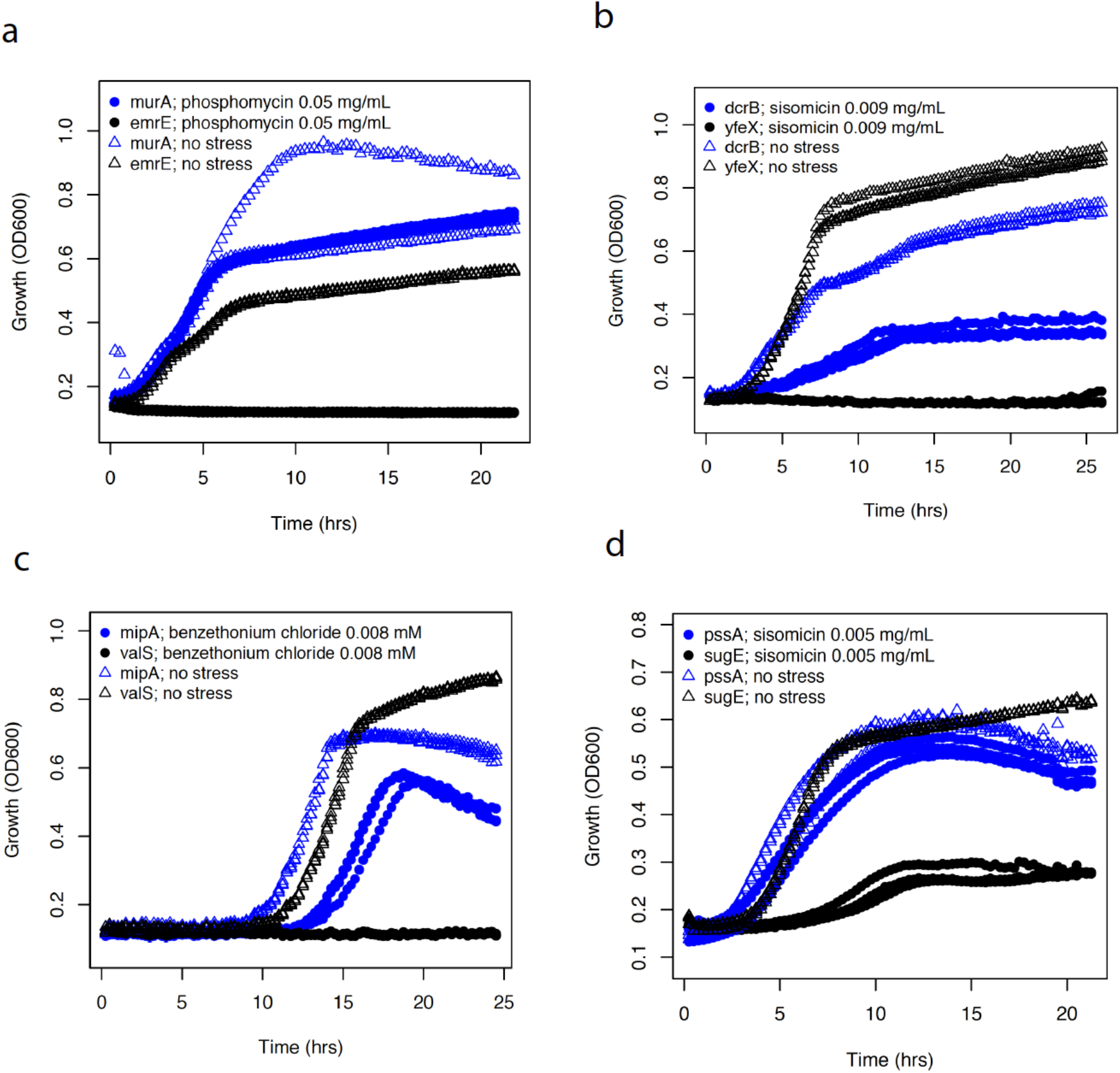
Additional validation growth curves for Dub-seq high scoring genes. (a) Growth of *E. coli* overexpressing *murA* under phosphomycin stress; *emrE* is a control. (b) Growth of *E. coli* overexpressing *dcrB* under sisomicin stress; *yfeX* is a control. (c) Growth of *E. coli* overexpressing *mipA* under benzethonium chloride stress; *valS* is used as a control. (d) Growth of *E. coli* overexpressing *pssA* under sisomicin stress; *sugE* is used as a control.

**Supplementary Fig. 6.**
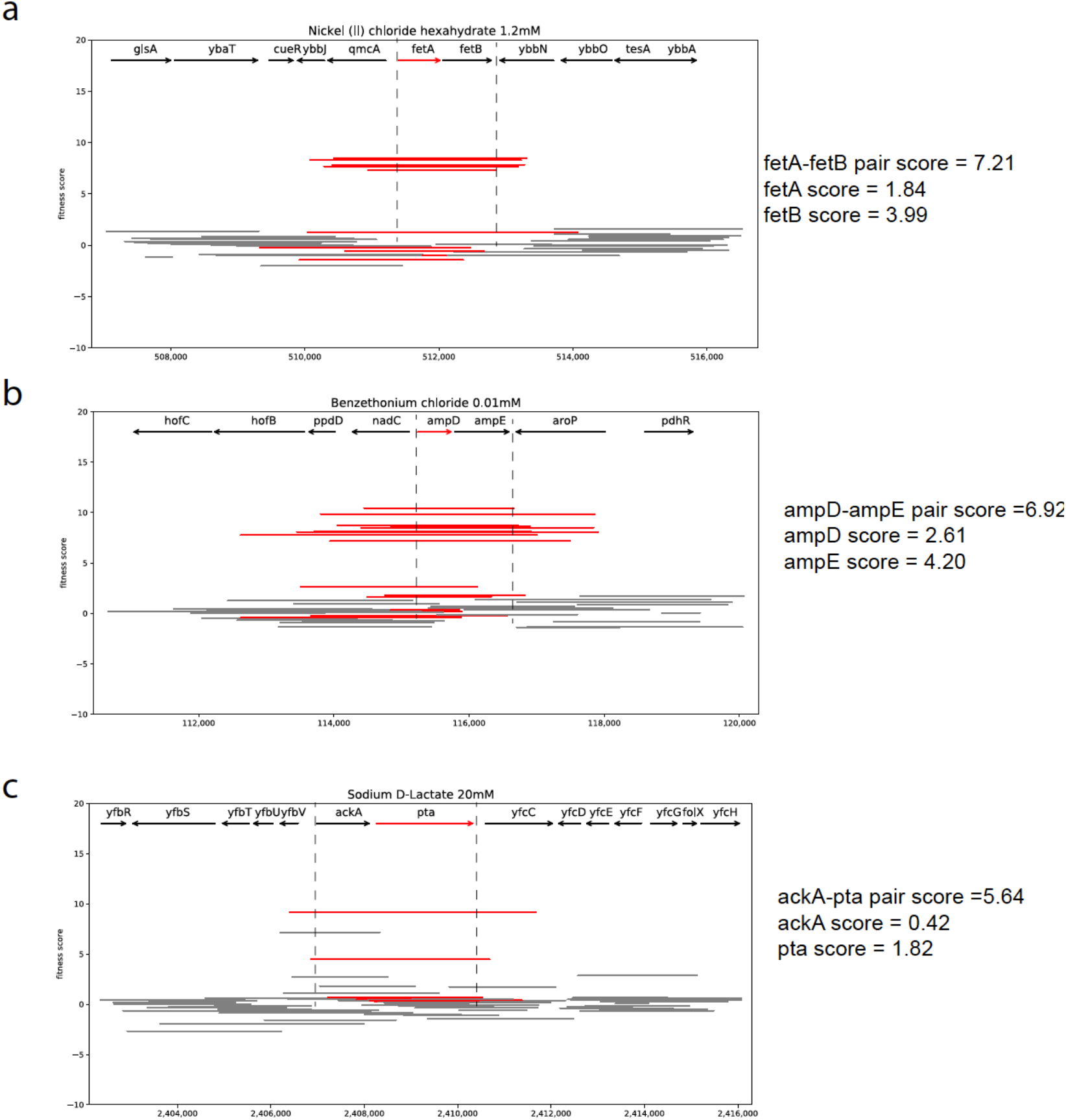
Dub-seq gene-pair fitness scores: Dub-seq fragment (strain) data (y-axis) for region surrounding gene-pair of interest (x-axis). The covered fragments are shown in red and partially covered gene-pair-neighborhood fragments are shown in gray. The regression scores each gene-pair of interest are shown next to each plot. Compare scores for (a) *fetA and fetB with fetA-fetB pair* with (b) *ampD and ampE, with ampD-ampE pair* and (c) *ackA and pta* with ackA-pta pair. We looked for the scores for fragments containing more than one gene that are significantly greater than the inferred sum of score of the constituent genes.

## Supplementary note

### Ridge, Lasso, and Elastic Net

The Ridge, Lasso, and Elastic Net regressions were implemented using the scikit-learn python library for machine learning. The regression was done on sparse representation of matrix A, without calculation of intercept since fragment scores were normalized (to set the median to zero). The regularization parameters were estimated using 3-fold cross validation (RidgeCV, LassoCV, and ElasticNetCV classes from the sklearn.linear_model package). The parameters were first estimated for each of 155 experiments, and then the parameters that deliver the highest R-square across all samples were selected as optimal.

The objective functions to be minimized and optimal regularization parameters for Ridge, Lasso, and Elastic Net are described below.

### Ridge

Ridge is *L*_2_ regularization with objective function:

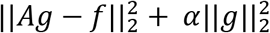

where ∝ controls the amount of regularization (shrinkage). The optimal *α* =1.0

### Lasso

Lasso is *L_t_* regularization with objective function:

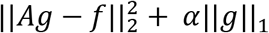

where *α* controls the amount of regularization (shrinkage) and variable selection. The optimal *α* =3.4

### Elastic Net

Elastic Net is regularization with linear combination of *L_t_* and *L_2_* terms and objective function:

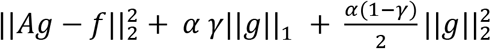

where *α* controls the amount of regularization and *α* defines the relative contribution of *L_t_* and *L_2_* terms/ The optimal parameters: *α* =3.6; *γ* =0.7

The regression analysis was run using optimal parameters and then manual inspection of regression results obtained from all three methods (Ridge, Elastic Net and LASSO) was performed for known gene-function associations. We observed that Ridge and Elastic Net with optimal parameters tends to significantly underestimate the fitness scores for causative genes that expected to have high positive or negative fitness scores. This underestimation is caused by shrinkage effect introduced by both regularization approaches. At the same time, the LASSO, when used with optimal parameters, seems to lack this problem and produces the most accurate scores across all three approaches. As an example, this is shown for *rcnA* gene (condition: 1.2 mM Nickel) scores calculated from Ridge, Elastic Net and LASSO approaches (**Supplementary Fig. 7a**). However, LASSO with optimal parameters still did not solve OLS over fitting problem completely, and still gave the unrealistic extreme positive and extreme negative scores for neighboring genes (for example, comparison of *rcnB* and *yehA*, condition: 1mM Cobalt, **Supplementary Fig. 7bc**). In comparison, NNLS had no regularization parameters, and we did not observe over fitting issues.

**Supplementary Fig. 7:**
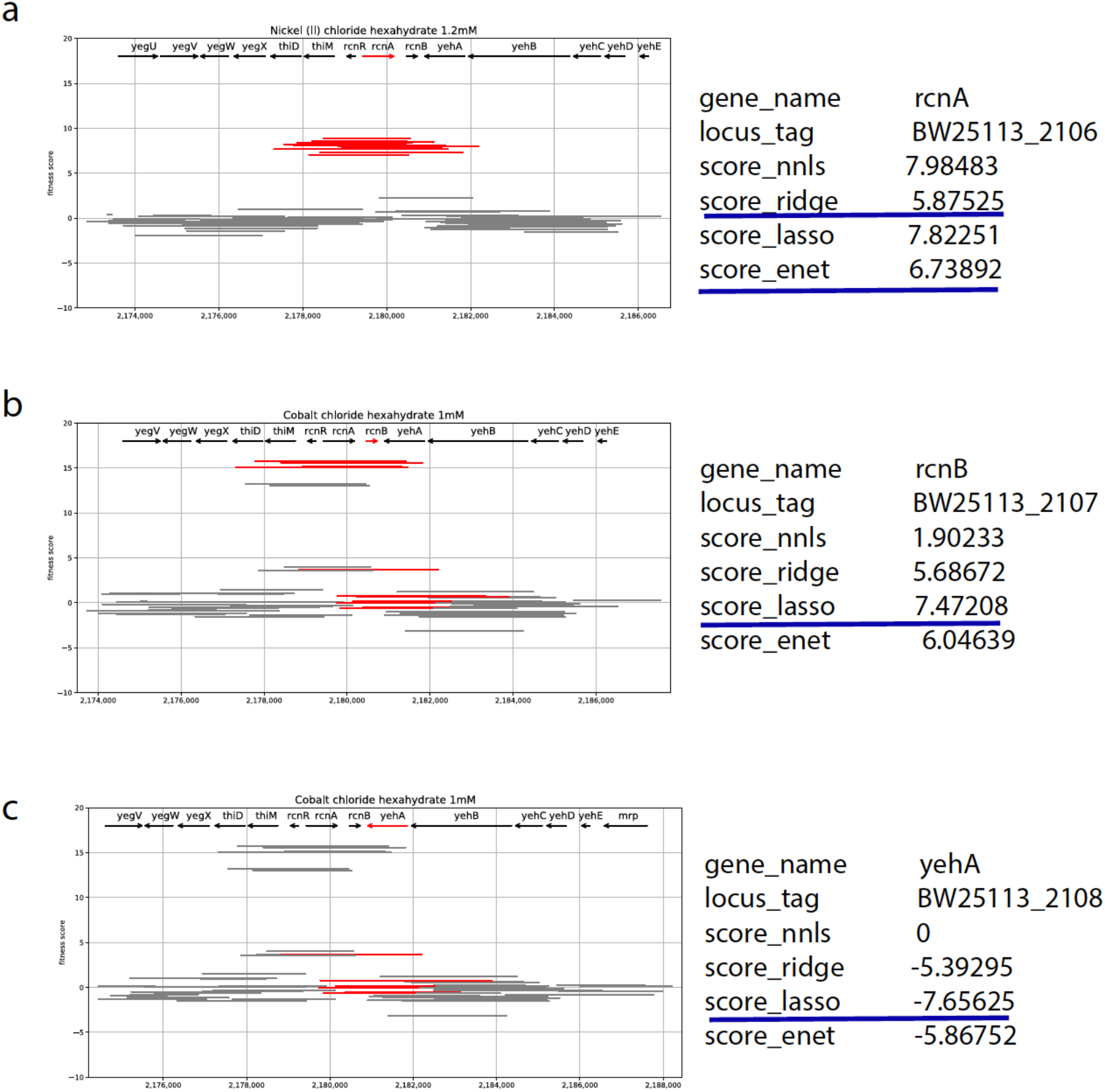
Gene score estimation approaches: Example gene scores for (a) *rcnA* (b) *rcnB* and (c) *yehA* showing data over fitting and shrinkage by ridge, lasso and elastic net regularization methods. Left, Dub-seq viewer for fragments covering a specific gene completely (red), compared to partially covering or gene-neighborhood fragments (gray). The gene scores estimated using different methods are shown on right. The gene scores highlighted in blue lines indicate issues of regularization methods (see Supplementary note).

## REFERENCES

1 Markowitz, V. M. et al. Ten years of maintaining and expanding a microbial genome and metagenome analysis system. Trends Microbiol 23, 7307–741, doi:10.1016/j.tim.2015.07.012 (2015).

2 Chang, Y. C. et al. COMBREX-DB: an experiment centered database of protein function: knowledge, predictions and knowledge gaps. Nucleic Acids Res 44, D330–335, doi:10.1093/nar/gkv1324 (2016).

3 Schnoes, A. M., Brown, S. D., Dodevski, I. & Babbitt, P. C. Annotation error in public databases: misannotation of molecular function in enzyme superfamilies. PLoS Comput Biol 5, e1000605, doi:10.1371/journal.pcbi.1000605 (2009).

4 Blaser, M. J. et al. Toward a Predictive Understanding of Earth's Microbiomes to Address 21st Century Challenges. MBio 7, doi:10.1128/mBio.00714-16 (2016).

5 Baba, T. et al. Construction of Escherichia coli K-12 in-frame, single-gene knockout mutants: the Keio collection. Mol Syst Biol 2, 2006 0008, doi:10.1038/msb4100050 (2006).

6 Koo, B. M. et al. Construction and Analysis of Two Genome-Scale Deletion Libraries for Bacillus subtilis. Cell Syst 4, 291–305 e297, doi:10.1016/j.cels.2016.12.013 (2017).

7 Giaever, G. & Nislow, C. The yeast deletion collection: a decade of functional genomics. Genetics 197, 451–465, doi:10.1534/genetics.114.161620 (2014).

8 Barker, C. A., Farha, M. A. & Brown, E. D. Chemical genomic approaches to study model microbes. Chem Biol 17, 624–632, doi:10.1016/j.chembiol.2010.05.010 (2010).

9 Brochado, A. R. & Typas, A. High-throughput approaches to understanding gene function and mapping network architecture in bacteria. Curr Opin Microbiol 16, 199–206, doi:10.1016/j.mib.2013.01.008 (2013).

10 Wang, H. H. et al. Programming cells by multiplex genome engineering and accelerated evolution. Nature 460, 894–898, doi:10.1038/nature08187 (2009).

11 Warner, J. R., Reeder, P. J., Karimpour-Fard, A., Woodruff, L. B. & Gill, R. T. Rapid profiling of a microbial genome using mixtures of barcoded oligonucleotides. Nat Biotechnol 28, 856–862, doi:10.1038/nbt.1653 (2010).

12 van Opijnen, T., Bodi, K. L. & Camilli, A. Tn-seq: high-throughput parallel sequencing for fitness and genetic interaction studies in microorganisms. Nat Methods 6, 767–772, doi:10.1038/nmeth.1377 (2009).

13 Wetmore, K. M. et al. Rapid quantification of mutant fitness in diverse bacteria by sequencing randomly bar-coded transposons. MBio 6, e00306–00315, doi:10.1128/mBio.00306-15 (2015).

14 Peters, J. M. et al. A Comprehensive, CRISPR-based Functional Analysis of Essential Genes in Bacteria. Cell 165, 1493–1506, doi:10.1016/j.cell.2016.05.003 (2016).

15 Price, M. N. et al. Mutant phenotypes for thousands of bacterial genes of unknown function. Nature 557, 503–509, doi:10.1038/s41586-018-0124-0 (2018).

16 Prelich, G. Gene overexpression: uses, mechanisms, and interpretation. Genetics 190, 841–854, doi:10.1534/genetics.111.136911 (2012).

17 Sandegren, L. & Andersson, D. I. Bacterial gene amplification: implications for the evolution of antibiotic resistance. Nat Rev Microbiol 7, 578–588, doi:10.1038/nrmicro2174 (2009).

18 Elliott, K. T., Cuff, L. E. & Neidle, E. L. Copy number change: evolving views on gene amplification. Future Microbiol 8, 887–899, doi:10.2217/fmb.13.53 (2013).

19 Rine, J., Hansen, W., Hardeman, E. & Davis, R. W. Targeted selection of recombinant clones through gene dosage effects. Proc Natl Acad Sci U S A 80, 6750–6754 (1983).

20 Ho, C. H. et al. A molecular barcoded yeast ORF library enables mode-of-action analysis of bioactive compounds. Nat Biotechnol 27, 369–377, doi:10.1038/nbt.1534 (2009).

21 Soo, V. W., Hanson-Manful, P. & Patrick, W. M. Artificial gene amplification reveals an abundance of promiscuous resistance determinants in Escherichia coli. Proc Natl Acad Sci US A 108, 1484–1489, doi:10.1073/pnas.1012108108 (2011).

22 Hoegler, K. J. & Hecht, M. H. Artificial Gene Amplification in Escherichia coli Reveals Numerous Determinants for Resistance to Metal Toxicity. J Mol Evol 86, 103–110, doi:10.1007/s00239-018-9830-3 (2018).

23 Qimron, U., Marintcheva, B., Tabor, S. & Richardson, C. C. Genomewide screens for Escherichia coli genes affecting growth of T7 bacteriophage. Proc Natl Acad Sci U S A 103, 19039–19044, doi:10.1073/pnas.0609428103 (2006).

24 Li, X. et al. Multicopy suppressors for novel antibacterial compounds reveal targets and drug efflux susceptibility. Chem Biol 11, 1423–1430, doi:10.1016/j.chembiol.2004.08.014 (2004).

25 Patrick, W. M., Quandt, E. M., Swartzlander, D. B. & Matsumura, I. Multicopy suppression underpins metabolic evolvability. Mol Biol Evol 24, 2716–2722, doi:10.1093/molbev/msm204 (2007).

26 Lynch, M. D., Warnecke, T. & Gill, R. T. SCALEs: multiscale analysis of library enrichment. Nat Methods 4, 87–93, doi:10.1038/nmeth946 (2007).

27 Nicolaou, S. A., Gaida, S. M. & Papoutsakis, E. T. Coexisting/Coexpressing Genomic Libraries (CoGeL) identify interactions among distantly located genetic loci for developing complex microbial phenotypes. Nucleic Acids Res 39, e152, doi:10.1093/nar/gkr817 (2011).

28 Dunlop, M. J. et al. Engineering microbial biofuel tolerance and export using efflux pumps. Mol Syst Biol 7, 487, doi:10.1038/msb.2011.21 (2011).

29 Kitagawa, M. et al. Complete set of ORF clones of Escherichia coli ASKA library (a complete set of E. coli K-12 ORF archive): unique resources for biological research. DNA Res 12, 291–299, doi:10.1093/dnares/dsi012 (2005).

30 Wang, H. H. et al. Genome-scale promoter engineering by coselection MAGE. Nat Methods 9, 591–593, doi:10.1038/nmeth.1971 (2012).

31 Freed, E. F. et al. Genome-Wide Tuning of Protein Expression Levels to Rapidly Engineer Microbial Traits. ACS Synth Biol 4, 1244–1253, doi:10.1021/acssynbio.5b00133 (2015).

32 Judson, N. & Mekalanos, J. J. TnAraOut, a transposon-based approach to identify and characterize essential bacterial genes. Nat Biotechnol 18, 740–745, doi:10.1038/77305 (2000).

33 Dong, C., Fontana, J., Patel, A., Carothers, J. M. & Zalatan, J. G. Synthetic CRISPR-Cas gene activators for transcriptional reprogramming in bacteria. Nat Commun 9, 2489, doi:10.1038/s41467-018-04901-6 (2018).

34 Leis, B., Angelov, A. & Liebl, W. Screening and expression of genes from metagenomes. Adv Appl Microbiol 83, 1–68, doi:10.1016/B978-0-12-407678-5.00001-5 (2013).

35 Ekkers, D. M., Cretoiu, M. S., Kielak, A. M. & Elsas, J. D. The great screen anomaly--a new frontier in product discovery through functional metagenomics. Appl Microbiol Biotechnol 93, 1005–1020, doi:10.1007/s00253-011-3804-3 (2012).

36 Uchiyama, T. & Miyazaki, K. Functional metagenomics for enzyme discovery: challenges to efficient screening. Curr Opin Biotechnol 20, 616–622, doi:10.1016/j.copbio.2009.09.010 (2009).

37 Sommer, M. O. A., Dantas, G. & Church, G. M. Functional characterization of the antibiotic resistance reservoir in the human microflora. Science 325, 1128–1131, doi:10.1126/science.1176950 (2009).

38 Munck, C. et al. Limited dissemination of the wastewater treatment plant core resistome. Nat Commun 6, 8452, doi:10.1038/ncomms9452 (2015).

39 Yaung, S. J. et al. Improving microbial fitness in the mammalian gut by in vivo temporal functional metagenomics. Molecular Systems Biology 11, 788–788, doi:10.15252/msb.20145866 (2015).

40 Gibson, M. K. et al. Developmental dynamics of the preterm infant gut microbiota and antibiotic resistome. Nat Microbiol 1, 16024, doi:10.1038/nmicrobiol.2016.242016).

41 Smith, A. M. et al. Quantitative phenotyping via deep barcode sequencing. Genome Res 19, 1836–1842, doi:10.1101/gr.093955.109 (2009).

42 Studier, F. W. & Moffatt, B. A. Use of bacteriophage T7 RNA polymerase to direct selective high-level expression of cloned genes. J Mol Biol 189, 113–130 (1986).

43 Sorek, R. et al. Genome-wide experimental determination of barriers to horizontal gene transfer. Science 318, 1449–1452, doi:10.1126/science.1147112 (2007).

44 Oh, J. et al. A universal TagModule collection for parallel genetic analysis of microorganisms. Nucleic Acids Res 38, e146, doi:10.1093/nar/gkq419 (2010).

45 Rodrigue, A., Effantin, G. & Mandrand-Berthelot, M. A. Identification of rcnA (yohM), a nickel and cobalt resistance gene in Escherichia coli. J Bacteriol 187, 2912–2916, doi:10.1128/JB.187.8.2912-2916.2005 (2005).

46 Romero, D. & Palacios, R. Gene amplification and genomic plasticity in prokaryotes. Annu Rev Genet 31, 91–111, doi:10.1146/annurev.genet.31.1.91 (1997).

47 Durfee, T. et al. The complete genome sequence of Escherichia coli DH10B: insights into the biology of a laboratory workhorse. J Bacteriol 190, 2597–2606, doi:10.1128/JB.01695-07 (2008).

48 Keseler, I. M. et al. The EcoCyc database: reflecting new knowledge about Escherichia coli K-12. Nucleic Acids Res 45, D543–D550, doi:10.1093/nar/gkw1003(2017).

49 Egler, M., Grosse, C., Grass, G. & Nies, D. H. Role of the extracytoplasmic function protein family sigma factor RpoE in metal resistance of Escherichia coli. J Bacteriol 187, 2297–2307, doi:10.1128/JB.187.7.2297-2307.2005 (2005).

50 Grabowicz, M. & Silhavy, T. J. Envelope Stress Responses: An Interconnected Safety Net. Trends Biochem Sci 42, 232–242, doi:10.1016/j.tibs.2016.10.002 (2017).

51 Nishino, K., Yamasaki, S., Hayashi-Nishino, M. & Yamaguchi, A. Effect of NlpE overproduction on multidrug resistance in Escherichia coli. Antimicrob Agents Chemother 54, 2239–2243, doi:10.1128/AAC.01677-09 (2010).

52 Guo, M. S. et al. MicL, a new sigmaE-dependent sRNA, combats envelope stress by repressing synthesis of Lpp, the major outer membrane lipoprotein. Genes Dev 28, 1620–1634, doi:10.1101/gad.243485.114 (2014).

53 Freeman, J. L., Persans, M. W., Nieman, K. & Salt, D. E. Nickel and cobalt resistance engineered in Escherichia coli by overexpression of serine acetyltransferase from the nickel hyperaccumulator plant Thlaspi goesingense. Appl Environ Microbiol 71, 8627–8633, doi:10.1128/AEM.71.12.8627-8633.2005 (2005).

54 Lim, B. et al. RNase III controls the degradation of corA mRNA in Escherichia coli. J Bacteriol 194, 2214–2220, doi:10.1128/JB.00099-12 (2012).

55 Couce, A. et al. Genomewide overexpression screen for fosfomycin resistance in Escherichia coli: MurA confers clinical resistance at low fitness cost. Antimicrob Agents Chemother 56, 2767–2769, doi:10.1128/AAC.06122-11 (2012).

56 Li, H., Zhang, D. F., Lin, X. M. & Peng, X. X. Outer membrane proteomics of kanamycin-resistant Escherichia coli identified MipA as a novel antibiotic resistance-related protein. FEMS Microbiol Lett 362, doi:10.1093/femsle/fnv074 (2015).

57 Silver, L. L. Fosfomycin: Mechanism and Resistance. Cold Spring Harb Perspect Med 7, doi:10.1101/cshperspect.a025262 (2017).

58 Nicolaou, S. A., Fast, A. G., Nakamaru-Ogiso, E. & Papoutsakis, E. T. Overexpression of fetA (ybbL) and fetB (ybbM), Encoding an Iron Exporter, Enhances Resistance to Oxidative Stress in Escherichia coli. Appl Environ Microbiol 79, 7210–7219, doi:10.1128/AEM.02322-13 (2013).

59 Hoon, S. et al. An integrated platform of genomic assays reveals small-molecule bioactivities. Nat Chem Biol 4, 498–506, doi:10.1038/nchembio.100 (2008).

60 Thompson, K. M., Rhodius, V. A. & Gottesman, S. SigmaE regulates and is regulated by a small RNA in Escherichia coli. J Bacteriol 189, 4243–4256, doi:10.1128/JB.00020-07 (2007).

61 Shuman, H. A. & Silhavy, T. J. The art and design of genetic screens: Escherichia coli. Nat Rev Genet 4, 419–431, doi:10.1038/nrg1087 (2003).

62 Grothe, S., Krogsrud, R. L., McClellan, D. J., Milner, J. L. & Wood, J. M. Proline transport and osmotic stress response in Escherichia coli K-12. J Bacteriol 166, 253–259 (1986).

63 Paradis-Bleau, C., Kritikos, G., Orlova, K., Typas, A. & Bernhardt, T. G. A genome-wide screen for bacterial envelope biogenesis mutants identifies a novel factor involved in cell wall precursor metabolism. PLoS Genet 10, e1004056, doi:10.1371/journal.pgen.1004056 (2014).

64 Neufeld, J. D. et al. Open resource metagenomics: a model for sharing metagenomic libraries. Stand Genomic Sci 5, 203–210, doi:10.4056/sigs.1974654 (2011).

65 Pal, C. et al. Metal Resistance and Its Association With Antibiotic Resistance. Adv Microb Physiol 70, 261–313, doi:10.1016/bs.ampbs.2017.02.001 (2017).

66 Gaida, S. M. et al. Expression of heterologous sigma factors enables functional screening of metagenomic and heterologous genomic libraries. Nat Commun 6, 7045, doi:10.1038/ncomms8045 (2015).

67 Ausubel, F. M. Short protocols in molecular biology : a compendium of methods from Current protocols in molecular biology. 5th edn, (Wiley, 2002).

68 Lee, T. S. et al. BglBrick vectors and datasheets: A synthetic biology platform for gene expression. J Biol Eng 5, 12, doi:10.1186/1754-1611-5-12 (2011).

69 Kovach, M. E., Phillips, R. W., Elzer, P. H., Roop, R. M., 2nd & Peterson, K. M. pBBR1MCS: a broad-host-range cloning vector. Biotechniques 16, 800–802 (1994).

70 Sambrook, J., Russell, D. W. & Sambrook, J. The condensed protocols from Molecular cloning : a laboratory manual. (Cold Spring Harbor Laboratory Press, 2006).

71 Lawson, C. L., Hanson, R. J. & Society for Industrial and Applied Mathematics. in Classics in applied mathematics 15 1 electronic text (xii, 337 p (Society for Industrial and Applied Mathematics (SIAM, 3600 Market Street, Floor 6, Philadelphia, PA 19104), Philadelphia, Pa., 1995).

